# Iron as a critical determinant for survival, oxidative stress, and infectivity of *Leishmania* parasites

**DOI:** 10.64898/2026.10.29.762707

**Authors:** Priya Priydarshni, Amir Zaidi, Parool Gupta, Vahab Ali

**Author notes:** To whom correspondence is to be addressed Dr. Vahab Ali.

## Abstract

Iron is an essential for all living organisms, including humans, and is present in RBCs & other iron-binding proteins that play crucial roles in oxygen transport, DNA synthesis & electron transport. Similarly, iron also plays a critical role in parasite biology. The unicellular *Leishmania* parasites scavenge essential iron inside host macrophages. We investigated the impact of iron on parasite growth, apoptosis, oxidative stress, and the expression of proteins involved in ROS, thiol metabolism, Fe-S clusters, and infectivity in macrophages. Our findings revealed that optimum iron concentrations promote parasite proliferation, whereas higher levels induce toxicity, leading to DNA fragmentation and caspase-like activity. Iron supplementation enhances parasite tolerance to oxidative stress by decreasing intracellular ROS and increasing total intracellular thiol content in a concentration-dependent manner. However, iron depletion increases ROS levels and decreases total thiol content in a concentration-dependent manner, which disrupts redox homeostasis. To support our finding, we further exposed these treated parasites to menadione, which supports that iron-supplemented cells can better survive under oxidative stress than the depleted cells. Furthermore, extra iron availability increases the expression of key metabolic proteins, Fe-S clusters (Nbp35, Frataxin, IScU) and thiol metabolism (TryS, TryR, CS, SAT, cTXN, cTXNPx), while under iron-depleted conditions these proteins were downregulated except frataxin. *In vitro* infectivity demonstrated that iron supplementation enhances parasite infectivity, while iron depletion impairs percentage infectivity. Our study demonstrates the roles of iron in the survival, redox regulation, and virulence of *L. donovani*, which can correlate with the role of cellular iron homeostasis in VL pathogenesis.

**Highlights:**

- Iron is crucial for cell growth, differentiation, apoptosis, oxidative stress, and expression of various proteins involved in thiol metabolism of *L. donovani*.
- Iron supplementation enhances parasite tolerance to oxidative stress decreasing intracellular ROS.
- Iron modulates the expression of key metabolic proteins, including Fe-S cluster assembly proteins (Nbp35, Frataxin, IScU) and thiol metabolism-related enzymes (TryS, TryR, CS, SAT, cTXN, cTXNPx).
- Iron supplementation enhances parasite infectivity, while iron depletion significantly reduced infectivity in macrophages.

## 1. Introduction

The visceral leishmaniasis (VL) is a vector-borne disease, also known as kala-azar in the Indian subcontinent, caused by the protozoan parasite *Leishmania donovani.* The disease is majorly transmitted to mammalian host by the bite of infected female *Phlebotomus* sandfly. This deadly illness poses significant health risks and can have severe consequences if not treated properly on time. If left untreated, this disease can be fatal in over 95% of cases. It primarily affects impoverished and immunocompromised populations in tropical and subtropical regions globally. The WHO estimated that approximately 0.05 to 0.09 million visceral leishmaniasis new cases emerges each year worldwide, with only 25% to 45% of these cases being officially reported as of year 2023 [1]. More than 90% of visceral leishmaniasis cases were reported in countries like, India, Ethiopia, Brazil, Kenya, Somalia, Sudan, and other countries with over 1 billion people at risk globally. However, approximately 70,000-1 million new cases of leishmaniasis are reported each year from nearly 90 endemic countries [2, 3]. VL is characterised by prolonged fever, hepatosplenomegaly, pancytopenia, and progressive immuno suppression, leading to high mortality if left untreated [4]. Anaemia has also been considered as a major haematological manifestation during VL. However, the cause of anaemia has been reported as multifactorial, such as hypersplenism and splenic sequestration, bone marrow suppression and ineffective erythropoiesis, inflammatory iron dysregulation, etc [5, 6]. Despite concerted control efforts, challenges such as drug resistance, inadequate healthcare infrastructure, and environmental changes sustaining vector populations continue to impede elimination [7]. The critical factor in *Leishmania’s* pathogenicity is its ability to exploit host iron metabolism to survive and replicate within the harsh phagolysosomal environment of host macrophages, where iron availability plays a pivotal role [8]. These unicellular protozoan parasites exhibit a unique survival strategy by residing within mammalian macrophages, where they harness host resources, mainly iron and heme, for essential metabolic processes [9]. Iron acquisition is particularly crucial for parasite proliferation, as *Leishmania* lacks a functional heme biosynthesis pathway and relies entirely on scavenging iron and heme from host macrophages [10]. Parasites have evolved sophisticated mechanisms to acquire, transport, and regulate iron, which is crucial for their survival within the host environment, while mitigating stress conditions and oxidative damage [11]. Iron is essential for many cellular possesses such as DNA synthesis, redox homeostasis, mitochondrial respiration, and redox balance, serving as a cofactor for enzymes like ribonucleoside diphosphate reductase and cytochrome oxidases [12]. *Leishmania* parasites has developed intricate mechanisms by following evolved pathways to acquire iron from host transferrin, ferritin, & heme, while concurrently suppressing macrophage iron-with holding defense [9]. For instance, *Leishmania* upregulates the expression of iron transporters such as LIT1 (*Leishmania* Iron Transporter 1) and exploits host transferrin receptor endocytosis to scavenge iron [13]. Additionally, the parasite’s reliance on heme oxygenase for heme catabolism underscores its adaptation to iron-scarce niches [14]. Studies have demonstrated that *Leishmania* species employ multiple iron uptake mechanisms, including involvement of several transmembrane protein like LFR (*Leishmania* Ferric Reductase), LIT (*Leishmania* Iron Transporter), LIR (*Leishmania* iron regularor) and heme-responsive pathways, to thrive within the iron-restricted environment of phagolysosomes [11, 13]. Apart from these adaptations, iron homeostasis in *Leishmania* is a double-edged sword; while sufficient amount is essential for growth, but excess iron can exacerbate oxidative stress through Fenton chemistry, necessitating tight regulatory control [15]. Understanding these mechanisms could unveil novel therapeutic targets, as current treatments for VL remain limited by toxicity, resistance, and high costs [16]. This study explores the intricate iron acquisition and utilization strategies of *Leishmania*, shedding light on their role in parasite virulence and persistence within the host.

Despite its importance, the precise effects of excess or insufficient iron on *Leishmania* survival and association of iron with anaemia remain unclear. The parasite completely relies on a specialized system, known as the *Leishmania* iron transporter (LIT), to regulate iron homeostasis within the parasitophorous vacuole [10]. Understanding these mechanisms not only sheds light on the intricate biology of parasitic infections but also highlights the delicate balance between host nutrition and pathogen survival strategies regarding infectious disease and further underscores the potential for targeted therapeutic interventions aimed at disrupting these essential pathways in combating leishmaniasis. Therefore, our work aims to explore the pivotal role of iron (Fe) transportation and utilization in *L. donovani*, focusing on their contribution to parasite survival, infectivity, and differentiation. We have also identified the one of the possible mechanisms which causes anaemia in VL and established the correlation of intracellular iron utilization and generation of anaemic condition in VL patients. By elucidating these pathways, the study seeks to uncover new insights into *Leishmania* pathogenesis and to potentially identify novel therapeutic strategies targeting iron metabolism for combating visceral leishmaniasis.

## 2. Material and method

### 2.1. Chemical and Reagent used in the study

All chemicals which were used in this study are of analytical grade. The parasites were cultivated using Gibco Medium 199 (M199) and Himedia Fetal Bovine Serum (FBS). THP-1 macrophages were grown in Sigma RPMI-1640 medium. All those chemicals and required reagents were sourced from reputed companies, including Sigma, Merck, Thermo-Fischer Scientific, USB, G-Biosciences, Takara, etc. The kits used for experiments are from Qiagen, Agilent, and G-Biosciences, etc.

### 2.2. Parasite culture and maintenance

The Wildtype (WT) promastigotes (MHOM/IN/1983/AG83) of *L. donovani* were cultivated in Medium 199 media mixed with 10% FBS (heat-inactivated) to make complete M199 medium, 25 mM HEPES buffer (pH 7.2), penicillin at 10 µg/ml, & streptomycin at 100 µg/ml. The promastigotes of *L. donovani* were maintained at 24±1 °C in Biochemical oxygen demand (BOD) incubator and further sub-cultured at every 4 to 5 days during the late logarithmic phase of parasite growth.

### 2.3. Promastigote growth in extra Iron supplementation and depletion

To analyse the effect of excess or insufficient iron (Fe) upon growth and survival of promastigote parasites of *L. donovani*, the parasites were cultivated in cM199 medium followed by addition of freshly prepared sterile ferrous sulphate heptahydrate (Fe_2_SO_4_ꞏH_2_O); with a final conc. of 0, 100, 250, 500, 1000, 2000 and 4000 µM, up to 5 days for iron supplementation as mentioned earlier. For iron depletion, WT promastigotes were cultivated in CM199 medium as mentioned in section 2.2 in the presence of freshly prepared sterile deferoxamine (DFO) at a final conc. of 0, 10, 20, 30, 40, and 50 µM to chelate iron from the medium and maintained up to 5 days. The WT Ag83 strain was used for iron supplementation and depletion [17]. A hemocytometer was used for cell counting after every 24 hrs post-incubation for parasites, and trypan blue exclusion method was followed to determine total cell number in the iron-supplemented and depleted conditions to evaluate the growth pattern.

### 2.4. Microscopy and Giemsa staining

Iron-supplemented and iron-depleted parasites were stained with Giemsa stain. To prepare giemsa stain, 670 µl of Giemsa Stain (Himedia) was taken and added to 30 ml of autoclaved distilled water. The culture was spread onto microscopic glass slides and allowed to air dry. After that, methanol was added onto a glass slide and again left to air dry for the fixation of cells. The prepared Giemsa stain was poured into a glass staining Coplin’s jar (Merck), and the slides were incubated inside for 45 mins at room temperature. After incubation, microscopic glass slides were washed with autoclaved distilled water for the removal of excess stain from the glass slide [18]. Microscopy was done at 100x resolution using immersion oil.

### 2.5. Monocyte cell culture and differentiation

The THP-1 cell lines were cultured in RPMI-1640 medium in a sterile culture flasks and supplemented with 10% FBS (heat-inactivated) and further maintained in a 5% CO_2_ incubator chamber. The complete medium was changed after every three days to insure optimal growth of THP-1 macrophages. After 3 days, 1×10^6^ cells/ml was seeded onto a coverslip in a 6-well tissue culture plate (TPP) in complete RPMI-1640 media added with 25 nM PMA and incubated for 72 hours in 5% CO_2_ incubator chamber at 37 °C before infection. During this phase, PMA induced THP-1 cell lines to differentiate into macrophages, enhancing their phagocytic characteristics and leading to morphological changes such as increased cell size and adherence to the coverslip. Non-adherent cells were removed using serum-free media. Subsequently, the PMA which acts as stimulus was removed by replacing the medium with fresh medium. The THP-1 cells were further incubated for 24 hrs to complete differentiation of macrophages [19]. The differentiated THP-1 cells were then infected with iron-supplemented and iron-depleted pre-treated stationary phase wild-type Ag83 strain of *L. donovani* parasites with ratio 1:10 (macrophage: parasite). The co-cultures were further incubated for 12 hours, followed by washing the cells from serum free media to remove non-internalised parasites. After the removal of the culture media, the attached macrophages were properly washed three times with using sterile Phosphate buffer saline (PBS). Plates containing the infected THP-1 macrophages were then further incubated for additional time points of 0, 24, and 48 hours under the same conditions with fresh complete media. Finally, the coverslips were fixed by using methanol and Giemsa stained and observed under an optical microscope (Olympus-BX41) with 100X magnification using immersion oil. A minimum of 500 cells, including both infected and non-infected cells were counted across randomly chosen microscopic fields per coverslip [20].

### 2.6. Preparation of soluble leishmanial lysate

The preparation of soluble leishmanial antigen (SLA) was performed as reported earlier [21, 22]. Briefly, 1x10^8^ cells/ml of iron-treated promastigotes of *L. donovani* was harvested from the stationary phase culture. The cultures were harvested, and the pellet was further dissolved in LLB (*Leishmania* lysis buffer) prepared by mixing 10 mg digitonin, 100 µl Protease inhibitor, 1 ml glycerol and PBS pH 7.4 up to 10 ml, respectively. The cell suspension was vortexed 4-5 times for 2 min each with a 10 min interval. The suspended parasites were further allowed to freeze-thaw 3 times in a liquid N_2_ cylinder for 10 minutes and kept in water bath for 5 min at 37 °C for thawing. The centrifugation of cell suspension was performed at 12000 rpm for 20 minutes at 4 °C. The collected supernatant containing cell lysate was collected and stored at -30 °C. The SLA concentration was measured by estimating protein through Bradford’s method using the CB protein assay kit, and absorbance was observed at 595 nm, followed by SDS-PAGE and immunoblotting.

### 2.7. Iron estimation

Total intracellular iron content in the prepared cell lysate from both iron-supplemented and iron-depleted cultures was estimated using the *O*-phenanthroline method of iron estimation. Briefly, 100-150 µg of total cell lysate (150 µl) proteins was taken and acidified using addition of 32N HCl (10 µl), then the sample was heated for 10 min at 80 °C. Then, the sample was mixed with 24 µl 3M sodium acetate, 12 µl 10% hydroxylamine hydrochloride, and 60 µl 0.1% *O-*phenanthroline. Then, samples were vortexed at room temperature (RT) for 30 minutes in the dark condition. Followed by incubation of reaction mixture at room temperature for 30 minute and at 512 nm absorbance was recorded using UV Spectrophotometer as described previously [23]. The Fe_2_SO_4_ (0-1000 µM) standard curve was prepared for the calculation of iron concentration.

### 2.8. Estimation of intracellular ROS

The reactive oxygen species was determined by following standardized method of our laboratory as described previously by our group [20, 24]. Basically, H_2_DCFDA or 2, 7,-dichloro-dihydrofluorescein diacetate (Sigma), a non-fluorecent form gets oxidized in presence of intracellular ROS and get converts in DCF or dichlorofluorescein, a green fluorescent form. The WT Ag83 strain of *L. donovani* was quantified in presence of both iron-supplemented and depleted conditions. Briefly, equal no. of cells (2×10^7^ parasites/well) in 100 µl cells suspension in PBS buffer were incubated with 20 µM H_2_DCFDA (2 µl) in dark for 30 min in a 96-well culture plate. The fluorescence spectrophotometer (Agilent Technologies) was used to measure the intensity of the produced fluorescence, by using excitation and emission wavelength at 504 nm and 525 nm, respectively. The fluorescence intensity was directly proportional to intracellular accumulation of ROS, and the result was expressed in relative fluorescence units (RFU). A blank containing 20 µM H2DCF-DA in PBS without parasites was used for comparison [25]. The experiment was conducted in duplicate.

### 2.9. Estimation of total reduced intracellular thiol

The estimation of total intracellular reduced thiol was determined as following our laboratory protocol [19, 20]. In short, 1x10^8^ parasite cells were pellet downed and further washed using PBS (pH 7.2), and then the cells were further resuspended in 600 µl of 25% TCA (Trichloroacetic acid). After precipitation, samples were incubated on ice for 10 minutes and the denatured protein including the extra-cellular debris were removed after centrifugation 12000 xg for 10 minutes at 4 °C. Then, the supernatant was collected and mixed with 3.6 µl of 0.6 mM DTNB (5-5, dithiobis-2-nitrobenzoic acid) also known as ellman’s reagent prepared in 200 mM sodium phosphate buffer (pH 8.0), respectively. The total intracellular thiol content was measured by UV-spectrophotometer at wavelength of 412 nm. The experiments were performed in three independent set and result data expressed as mean ± standard error of mean.

### 2.10. MTT Viability Assay

To determine the cell viability of *L. donovani* cultured in iron-supplemented and iron-depleted conditions, the MTT assay was used as described earlier standardized in our laboratory [26]. Exponentially growing 1x10^6^ cells/ml *Leishmania* parasites were cultivated in cM199. Briefly, 1x10^6^ promastigotes per ml were aliquoted in a 96-well culture plate which was further treated with increasing conc. of both iron supplementation and iron depletion by using deferoxamine. After that, MTT reagent added with final conc. of 400 µg/ml & followed by 3 hours incubation at 25 °C. The resulting crystals of formazan were solubilized using 200 µl MTT solubilization buffer. The ELISA plate reader (Bio-Rad) was used to measure absorbance at 570 nm [27, 28]. Experiments were performed thrice in three independent sets and statistical analysis performed by using GraphPad Prism 5.0 software.

### 2.11. Immunoblotting analysis

WT Ag83 strain of *L. donovani* parasite culture was cultured in both iron-supplemented and iron-depleted media conditions. A total of 30 µg of protein was estimated and loaded in SDS-PAGE gel. The protein estimation was performed by using CB protein assay and for protein standard curve preparation, 1.0 mg/ml BSA (Stock) was used as standard protein as reported earlier [22, 25, 29]. The promastigote total cell lysate was resolved on 10%, 12% and 15% (depending upon the size of the protein). The SDS-PAGE was performed and followed by western blotting, where 0.22µ PVDF membrane (GE healthcare) was electroblotted. Then, the protein transferred to PVDF membrane incubated with 5% skimmed milk prepared in TNT buffer for blocking and kept for an hour on a shaker at room temperature followed by probing using polyclonal antibody. For, TNT buffer preparation, 100 mM Tris-Cl of pH=8.0, 0.05% Tween-20 solution and 150 mM of NaCl was combined. The polyclonal antibodies also called as primary antibodies were used in different dilutions as mentioned: anti-LdTryS, anti-LdActin, anti-LdCS (1:3000), anti-LdSAT (1:2000), anti-Nbp35 (1:1500), anti-LdFrataxin, anti-LdcTXN, anti-TryR, anti-LdcTXNPx & anti-LdIScU (1:1000). These antibodies were generated in our lab against purified proteins in rabbit. The mentioned dilutions of various antibodies were used in the blocking buffer and incubated on a shaker for 1 hour at RT. After that, the PVDF membrane was washed by using TNT buffer thrice at an interval of every 10 minutes. The membrane was further incubated in secondary-Ab conjugated with AP *i.e.* alkaline phosphatase (goat anti-rabbit IgG) with 1:5000 dilutions and kept on shaker for 1 hr at RT as mentioned earlier [30]. Again, PVDF membrane washed using TNT buffer as mentioned above & PVDF membrane blot was developed using 5 µl of both BCIP/NBT substrate solution (Sigma-Aldrich) added in 5 ml Alkaline phosphatase buffer. ImageJ software was used for densitometric analysis to analyse the difference in band intensities, and fold change in the protein expression was analysed.

### 2.12. Caspase-like Activity

Caspase-like activity in WT Ag83 strain of *L. donovani* was assayed by using the kit Generic Caspase activity assay kit (Abcam) as used earlier [31]. This assay was performed by following manufacture’s guideline and the given protocol was followed properly. Briefly, WT parasites from both iron-supplemented and depleted cultures were used to evaluate apoptosis. 100 µL (1x10^6^ cells/ml) of parasite culture was taken from each sample in 1.5 ml MCT, and 1 µl of 500X TF2-VAD-FMK was mixed in the tube. Then, the parasitic cells were kept in dark condition at 37 °C in 5% CO_2_ incubator for four hrs. After incubation, cells were washed with PBS buffer for removal of extra unbounded reagent and then the cells were dissolved in 500 µl of assay buffer. We have identified fluorescence intensity by using spectrofluorometer from Agilent at the wavelength 490 nm (excitation) and 525 nm (emission).

### 2.13. Menadione induced oxidative stress

The Fe-supplemented and depleted WT promastigotes were further exposed to menadione to investigate the limit of oxidative stress tolerance. For *in vitro* oxidative stress generation, 1 × 10^6^ cells/ml iron-treated promastigotes of *L. donovani* parasites were cultured in 2.5 µM and 5.0 µM in a 96-well culture plate for a time period of 0-18 hrs [22, 23, 25]. Further, cell counted microscopically through hemocytometer by trypan blue exclusion method for 0-18 hrs with intervals of 3 hrs. At last, the percentage cell viability was calculated, and graph was plotted by GraphPad Prism (version 5.0).

### 2.14. DNA Fragmentation

DNA was isolated manually from both iron-supplemented and depleted WT parasites by the Phenol-Chloroform extraction method, as reported earlier [32]. DNA was quantified using the nanodrop (Genetix). 100 ng DNA from each concentration of iron-supplemented and depleted parasites was calculated and loaded in 1% agarose gel. DNA fragmentation was observed on gel and bands were visualised using the UV Gel documentation facility (Bio-Rad).

### 2.15. Statistical assessment

We have performed statistical analysis of all experiments and data has been represented as mean ± standard error of all experiments conducted in duplicate or triplicates. We have used GraphPad Prism 5.0 for data analysis. We have used one-tailed student t-test for smaller datasets and ANOVA for complex data groups. The difference between control and experimental group if less than 0.5 then it will be considered as significant. Significant differences has been represented as p value < 0.001 higher significant, p value < 0.01 significant, p value < 0.05 less significant and ns represented as non-significant values.

## 3. Results

### 3.1. Iron is necessary for growth and survival of *Leishmania* parasites

Many cellular biochemical functions, structural activities, and metabolic pathways including gene expression and regulation as well as cell proliferation depend completely on iron. Iron has reported to influence the cell fate i.e. the level of iron determines if the cell will proliferate or die [33–35]. The impact of external iron supplementation using Ferrous sulphate heptahydrate, *L. donovani* parasite development and survivability were examined in wild-type (WT) Ag83 promastigote strain cultured in cMI99 medium [36] **(Fig. 1A)**. WT strain growth pattern revealed that the highest cell proliferation occurred at 500 µM iron concentration on the 5th day, surpassing all other iron concentrations. Growth was consistently enhanced at increasing concentration of iron as compared to the control up to 5th day. However, by the 6th day, growth began to decline among all Fe concentrations, ranging from 100 µM to 4000 µM. Specifically, up to 2000 µM Fe concentration, the growth pattern of the parasites was higher depicted a modest increase in cell growth in comparison to the control. Beyond 2000 µM, iron concentrations were found to be toxic, significantly inhibiting the growth of the promastigotes **(Fig. 1B)**. The observed growth pattern of the WT strain in response to varying iron concentrations suggests a biphasic effect of iron on *Leishmania* parasites. The enhancement of cell proliferation at 500 µM iron on the 5th day indicates that iron is a critical factor for optimal growth within a certain range of concentration. Iron supplementation also enhanced promastigote proliferation (around 1.8-fold higher density at 5^th^ day, p < 0.001***). Although, the promastigote growth pattern was similar at both 1000 µM and 2000 µM which shows cell manages to survive and proliferate at 2000 µM iron concentration. Our result confers the role of iron as an essential micronutrient in cellular metabolism and cell differentiation. However, the transition from growth enhancement to inhibition above 2000 µM highlights a critical threshold beyond which iron concentration becomes detrimental. This dual effect underscores the importance of maintaining iron homeostasis in parasitic growth and could have implications for understanding how *Leishmania* parasites adapt to varying iron environments in the host. Our result confirmed that parasite cells showed better growth and survival in optimum iron supplemented condition while dense iron deposit or iron loaded conditions hampered cell growth and differentiation. This illustrated that iron supplementation elevated intacellular iron levels & were effectively working. However, we concluded that iron had a direct effect on parasite physiology, as growth of the promastigotes changed with different iron level.

**Fig 1.**
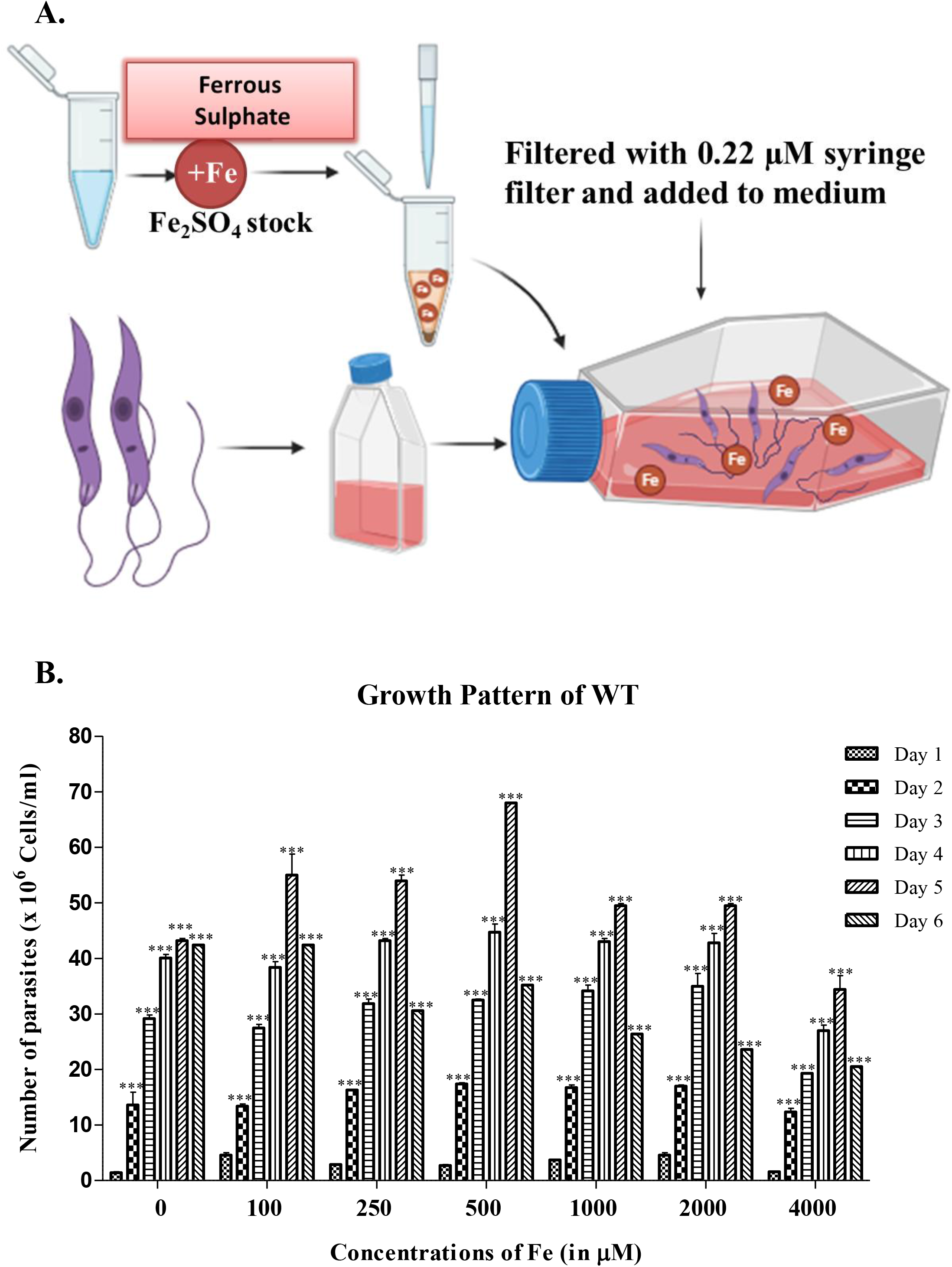
Growth pattern of iron supplemented promastigotes of *L. donovani.* **A.** The schematic illustration show**s** culture set-up condition for Fe supplementation. **B.** This graph represents the growth pattern of WT Ag83 promastigotes of *L. donovani* parasites (1×10^6^ cells/ml) cultured in increasing iron concentration 0-4000 µM Fe_2_SO_4_ for iron supplementation for 5 days in M199 medium with 10% fetal bovine serum. Cells were counted by trypan blue exclusion method using haemocytometer at every 24 hrs time interval for 6 days. The data represent mean ± SEM from three independent triplicate experiment indicates ns for non-significant, p < 0.05* and p < 0.001*** analysed grouped datasets by two-way ANOVA.

### 3.2. Iron depletion arrests promastigote proliferation

We further evaluated the effect of external iron depletion by using an extracellular iron chelator deferoxamine (DFO) in *L. donovani* promastigotes. The WT Ag83 strain was cultivated in cM199 medium by adding deferoxamine to analyse the effect of Fe deficiency upon *Leishmania* parasite growth [37]. Promastigotes were co-cultured in the presence of DFO at different concentrations from 0, 10, 20, 30, 40 & 50 µM for up to 6 days **(Fig. 2A),** and cell density was measured by counting the cells at every 24-hour interval by using hemocytometer after trypan blue staining. Cell viability decreased due to iron depletion, which was directly proportional to DFO concentration and the duration of exposure of DFO. At 4^th^ day, our result showed iron depletion suppressed growth reduction by 14.28% at 10 µM, 18.14% at 20 µM, 24.21% at 30 µM, 38.57% at 40 µM and 57.69% at 50 µM, p < 0.001*** up to the 4^th^ day. However, when intracellular Fe was depleted by 50 µM DFO, there was a significant cell death **(Fig. 2B)**. There was also a very significant decrease in the growth curve in higher iron chelation at 50 µM DFO as compared to control **(Fig. 3A)**. Our result depicts that treatment with DFO (50 µM) for one day led to a 46.22% decrease (p < 0.1**) and five days led to an 80.19% decrease (p < 0.001***) in intracellular iron. The lower physiological concentration 10 µM deferoxamine (9.42% iron chelation) also has an impact on parasite survival reduction **(Table 1)**. We observed that WT parasites are dependent on iron, and in iron-deficient conditions, they show poor physiology, inhibiting parasite survival **(Fig. 3B)**. It is evident that changes in intracellular iron level had a significant impact on *Leishmania* parasite survivability and cell differentiation.

**Fig 2.**
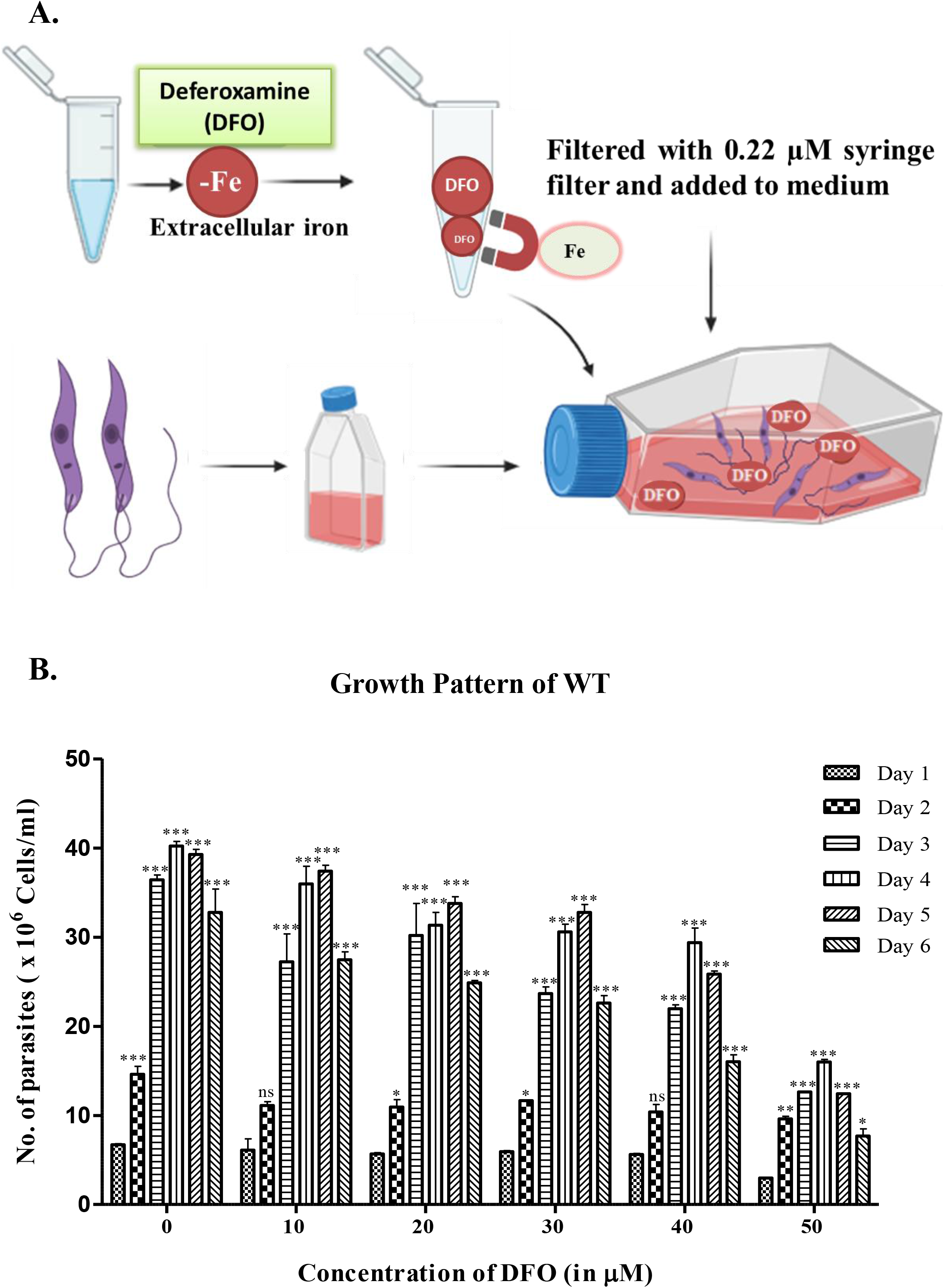
Growth pattern of iron depleted promastigotes of *L. donovani.* **A.** The schematic illustration show**s** culture set-up condition for Fe depletion. **B.** This graph represents the growth pattern of WT Ag83 promastigotes of *L. donovani* parasites (1×10^6^ cells/ml) cultured in increasing iron chelation 0-50 µM DFO for iron depletion for 5 days in M199 medium with 10% fetal bovine serum. Similarly, cells were counted by trypan blue exclusion method using haemocytometer at every 24 hrs time interval for 6 days. The plotted data represent mean ± SEM from three independent triplicate experiment indicates ns for non-significant, p < 0.05* and p < 0.001*** analysed grouped datasets by two-way ANOVA.

**Fig 3.**
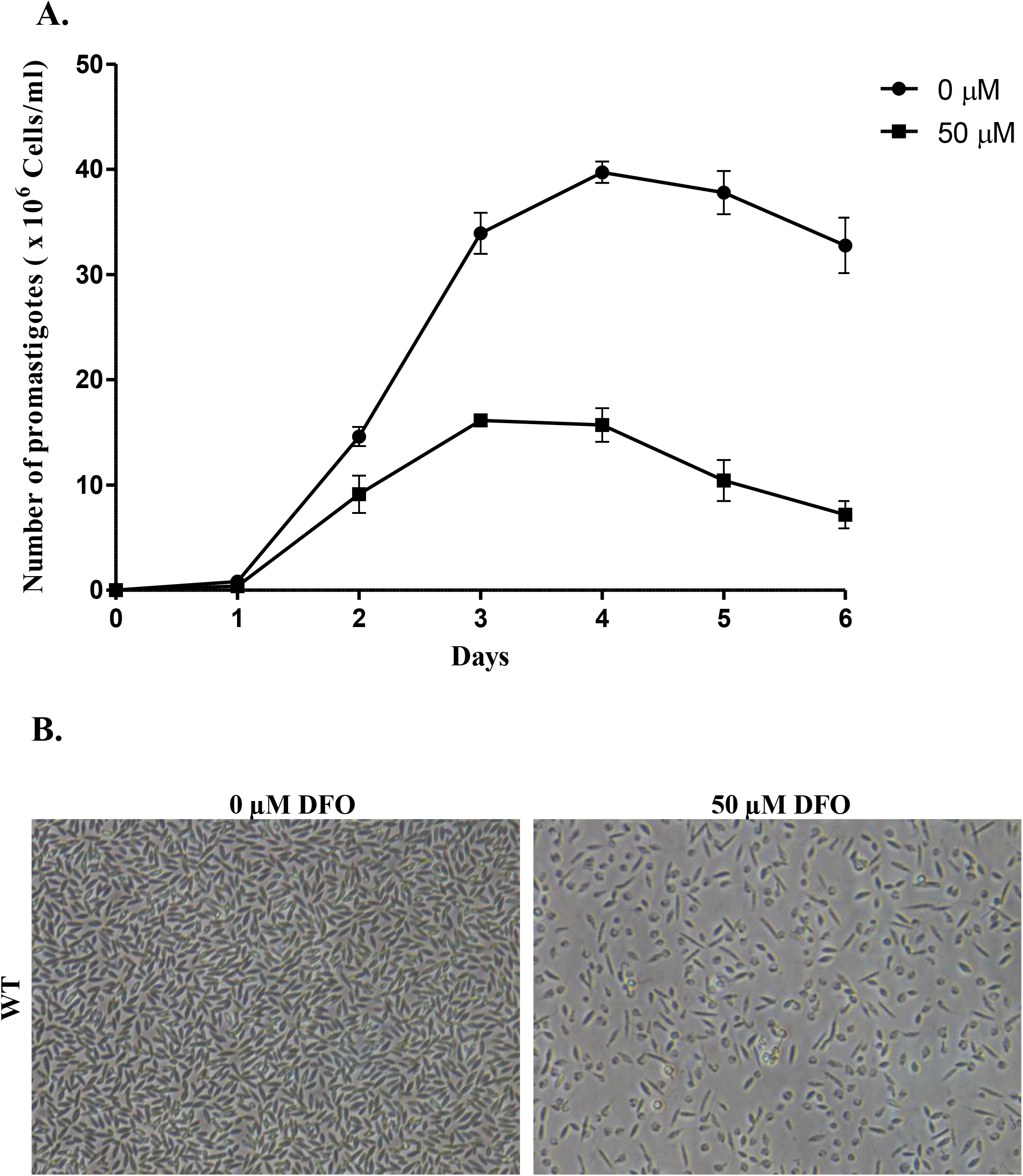
Iron chelation lowers the cell growth of WT parasites of *L. donovani*. **A.** 1×10^6^ parasites/ml were cultured in M199 medium in the presence of 50 µM DFO in comparison to control for 6 days. The cell counting was performed at every 24-hour interval. Growth curve was plotted using GraphPad software 5.0. **B.** Cell morphology was observed at 40x magnification and images were taken using an inverted microscopic of WT promastigotes parasites cultured in 0 µM DFO and 50µM DFO iron depleted conditions. Scale bar represents 10 µm as distance.

**Table 1.**
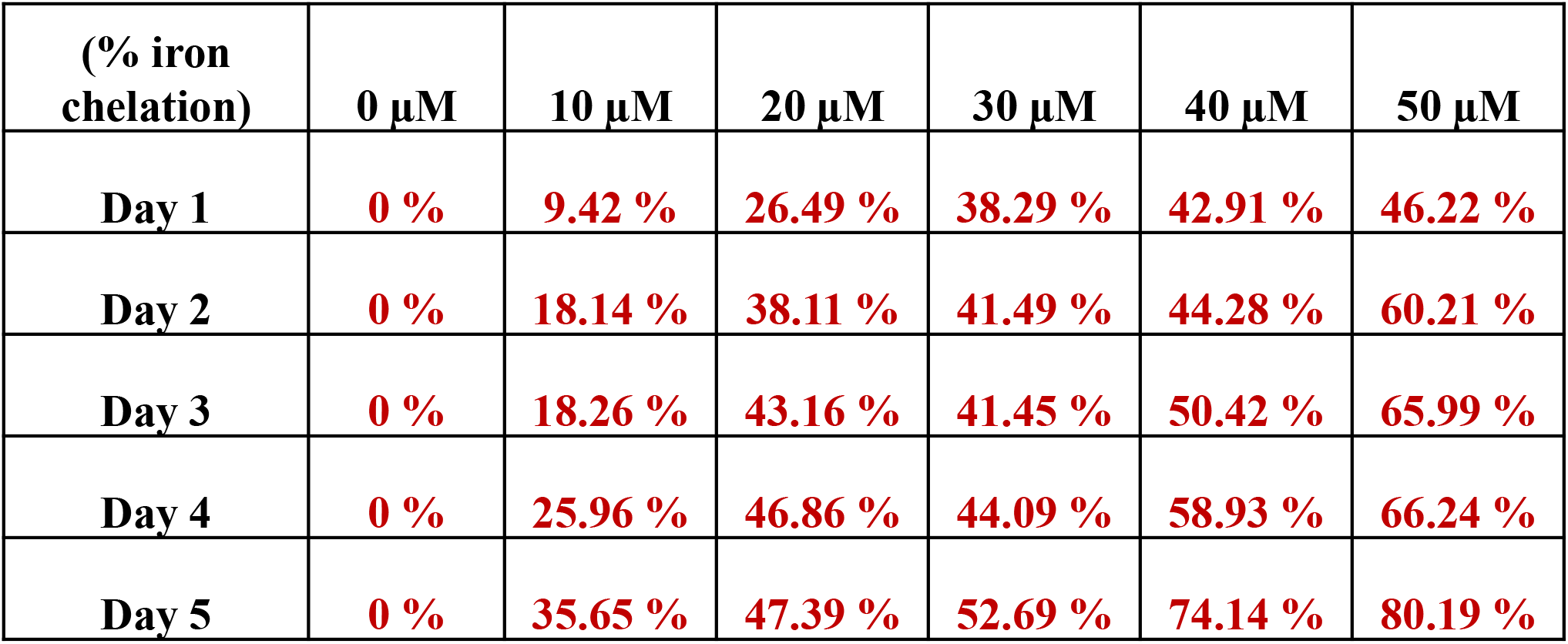
This table depicts average value of percentage iron chelation occurred due to deferoxamine (an extracellular iron chelator) treatment for iron depletion in WT parasites foe 5 days. Logarithmic phase *L. donovani* promastigotes (1×10^6^ cells/ml) were cultured in CM199 medium with the addition of increasing concentrations of Deferoxamine (0, 10, 20, 30, 40 & 50 µM) for iron depletion. The experiment was performed in triplicate, and Iron chelation was estimated.

### 3.3. Confirmation of iron supplementation and depletion

To determine whether the availability of iron hampered the cell growth, we estimated intracellular iron in the promastigotes cultured under iron-supplemented and iron-depleted conditions. We observed that total intracellular iron was significantly increased with higher iron supplementation **(Fig. 4A)** and decreased sequentially from 10-50 µM DFO under iron depletion **(Fig. 4B)**. Promastigotes cultured in iron-containing medium (1000 µM ferrous sulphate heptahydrate) showed around 1.3-fold increase in intracellular iron content compared to control on the 4^th^ day **(Fig. 4A)**. Fe supplementation confirmed elevated iron levels (9.17 µM Fe/mg at 1^st^ day, 4000 µM) in supplemented parasites vs untreated control. Treatment of parasites with deferoxamine at 10 µM further led to around a 10% decrease in the intracellular labile iron pool. Iron estimation showed around a 3-fold reduction in total intracellular iron in 50 µM DFO-treated cells on the 1^st^ day compared to control. We found that in iron-depleted conditions, iron was depleted with time and in concentration-dependent manner. With increasing conc. of DFO, iron level decreased inside parasites, and with increasing concentration of iron supplementation, iron level was increased in the parasites, demonstrating that iron supplementation and iron depletion were successful and affected iron acquisition and transportation in the parasitic cells. Our result confirms successful iron modulation in *Leishmania* parasites, as measured through Fe estimation, via iron supplementation (Fe_2_SO_4_ꞏ7H_2_O) and iron depletion (DFO).

**Fig 4.**
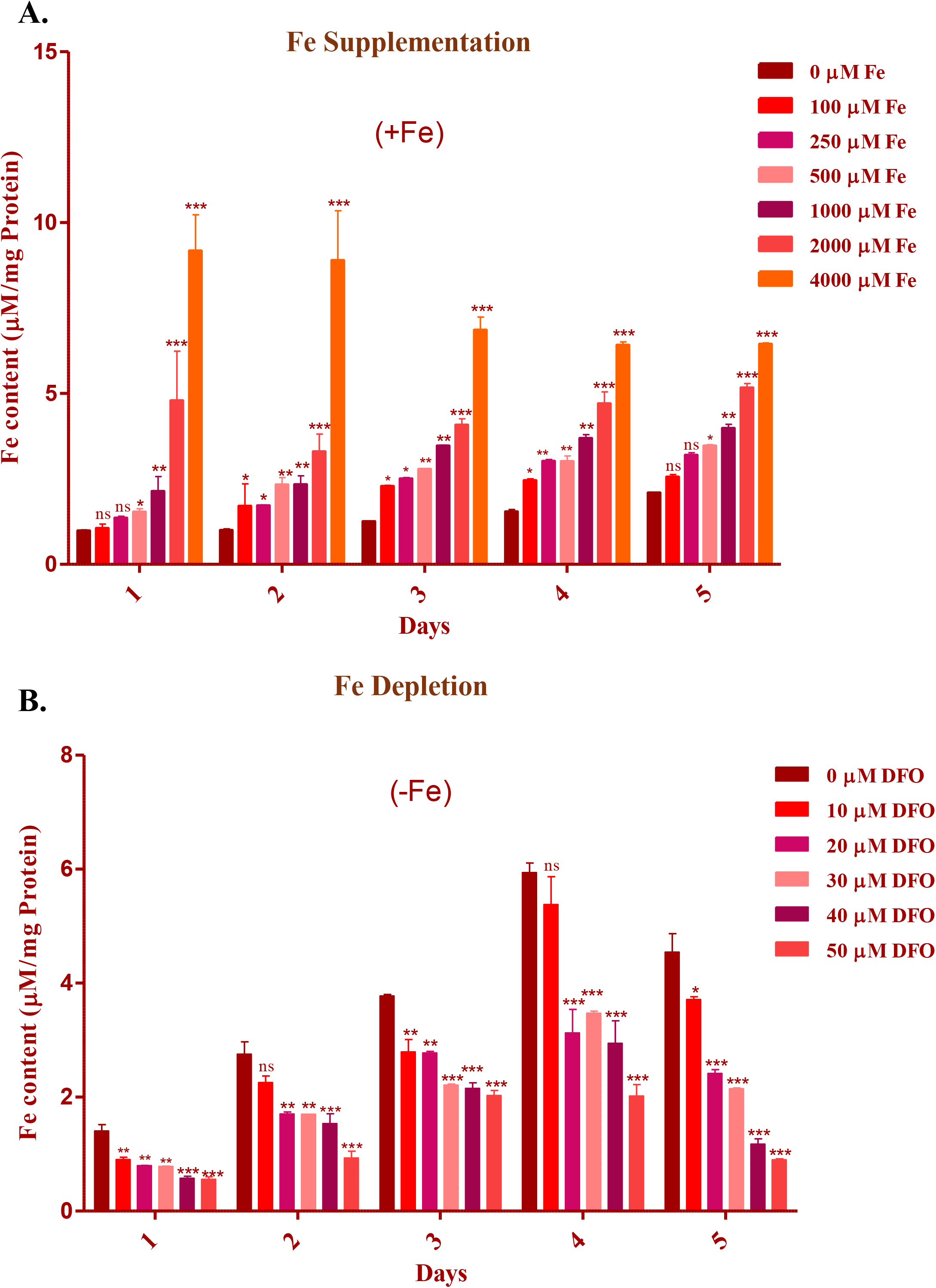
Intracellular iron estimation in *L. donovani* promastigotes after iron supplementation and iron depletion conditions. For iron estimation, *L. donovani* promastigotes were cultured in both iron supplemented (0-4000 µM Fe_2_SO_4_) and iron depleted (0-50 µM deferoxamine) media condition for five days and Fe estimation was performed at every 24-hr time interval for 5 days for both **(A.)** iron supplemented condition and **(B.)** iron depleted condition using *o*-phenanthroline method as mentioned in methodology section in triplicates. The untreated cells are taken as control and shown as 0 µM for both in iron-supplemented and iron depleted conditions. Each data point represents the mean ± SEM from three independent experiments performed in triplicate and Fe content is denoted as µM/mg protein of total cell lysate. Statistical analysis shows ns for non-significant, *p < 0.05, **p < 0.01 and ***p < 0.001 by using two-way ANOVA followed by Bonferroni test.

### 3.4. Fe supplementation and depletion effects parasites morphology

Distinct morphological changes were observed after both iron-supplemented and iron-depleted conditions. Parasites treated with lower iron concentration (0-1000 µM) for 5 days were substantially increased in cell size, became elongated shape with longer flagella as compared to untreated control, but higher iron treatment (2000 & 4000 µM) led to toxicity inside the cell and resulted in shrinkage, disoriented cell shape, size, and shorter flagella **(Fig. 5A)**. Conversely, treatment with DFO (0-50 µM) for 5 days led to a substantial reduction in size, cell shape was shrunk, and flagella shortened with an ovoid cell shape compared to control **(Fig. 5B)**. However, we observed no such significant difference in parasite morphology after incubating parasites at lower concentrations of Fe_2_SO_4_ (100 & 250 µM) and DFO (10-20 µM) after five days. Thus, our result depicts that iron-rich or poor condition certainly has a visible impact on *Leishmania* promastigotes morphology.

**Fig 5.**
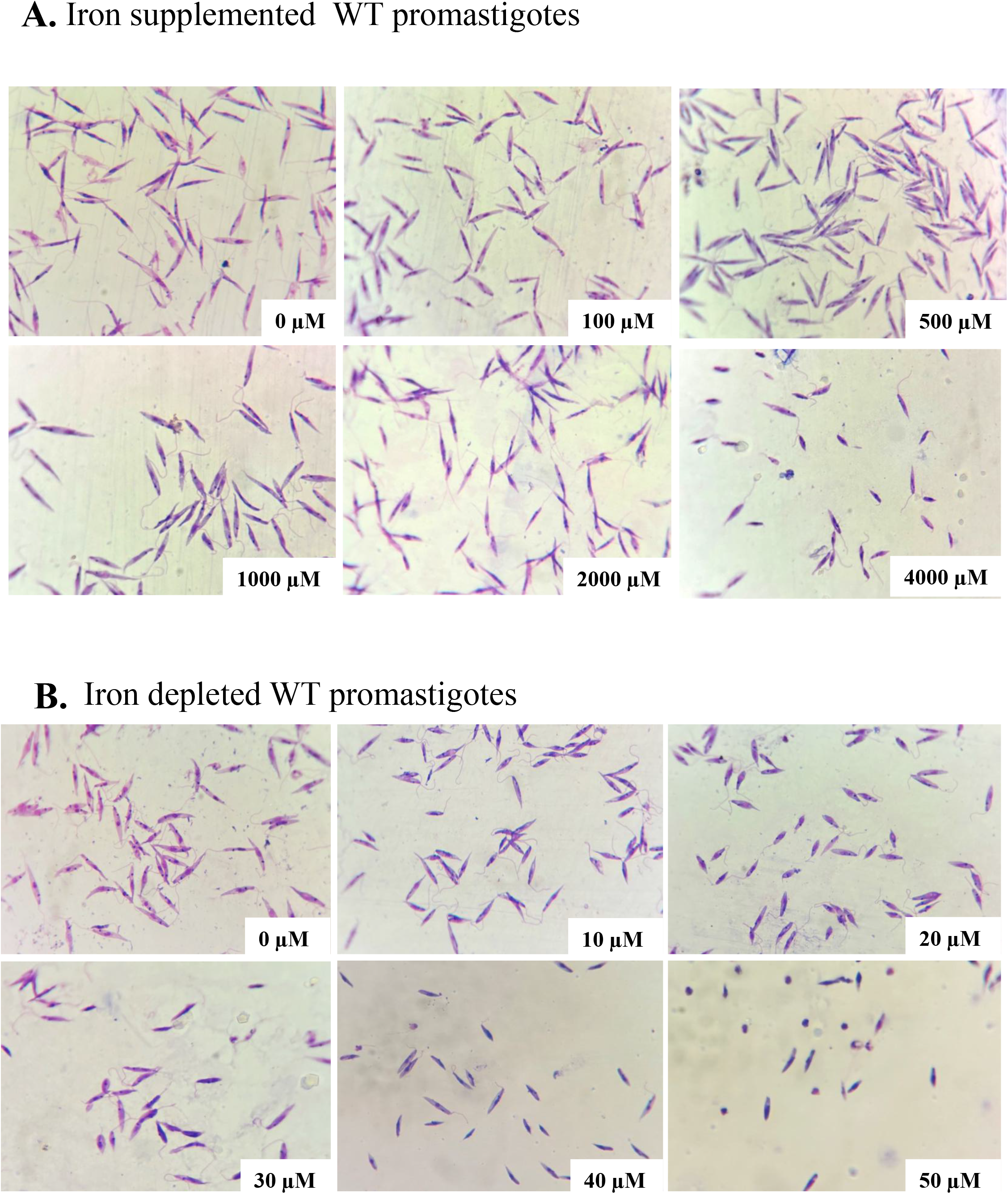
Morphological analysis of iron treated WT promastigotes of *L. donovani*. The parasites were cultured in CM199 medium and treated with **A.**0, 100, 500, 1000, 2000 & 4000 µM Fe_2_SO_4_ and **B.** 0, 10, 20, 30, 40 & 50 µM for 5 days and Giemsa staining was performed as mentioned in the method and material section. Untreated cells are shown as 0 µM in both respective conditions. The image was taken at 100x resolution using compound microscopy.

### 3.5. Iron affects *L. donovani* promastigotes viability

We treated promastigotes of *L. donovani* with increasing conc. of both iron supplementation (0-4000 µM_ Fe_2_SO_4_ꞏ7H_2_O) and depletion (0-50 DFO) up to 5 days, and MTT assay was used to measure cell viability of treated parasites. Our findings demonstrated that the cytotoxicity of promastigotes grown in iron supplemented and iron-depleted medium condition was both time- and concentration-dependent. We detected cell viability around ̴ 1.2-fold increase at 100 µM, ̴ 1.5-fold increase at 250 µM, ̴ 1.7-fold increase at 500 µM, ̴ 1.3-fold increase at 1000 µM, ̴ 1.3-fold decrease at 2000 µM after four days in iron supplementation, respectively **(Fig. 6A).** Our result showed a maximum of around 1.7-fold increases with p < 0.001*** in cell viability after 5 days in response to 500 µM iron supplementation **(Fig. 6A)**. Cell viability at the higher iron concentration 4000 µM was not included because the cell count at this concentration was insufficient for reliable analysis. We further evaluated the *L. donovani* parasites cell viability in presence of DFO which showed ̴ 1.2-fold decrease at 10 µM, ̴ 1.7-fold decrease at 20 µM, ̴ 2.3-fold decrease at 30 µM, ̴ 4.2-fold decrease at 40 µM, and ̴ 5.8-fold decrease at 50 µM in compared to control dataset **(Fig. 6B)**. Since, iron is essential for the *Leishmania* parasite’s survival [37, 38], growth pattern, and cell viability are clearly hampered due to iron-sufficient and deficient conditions.

**Fig 6.**
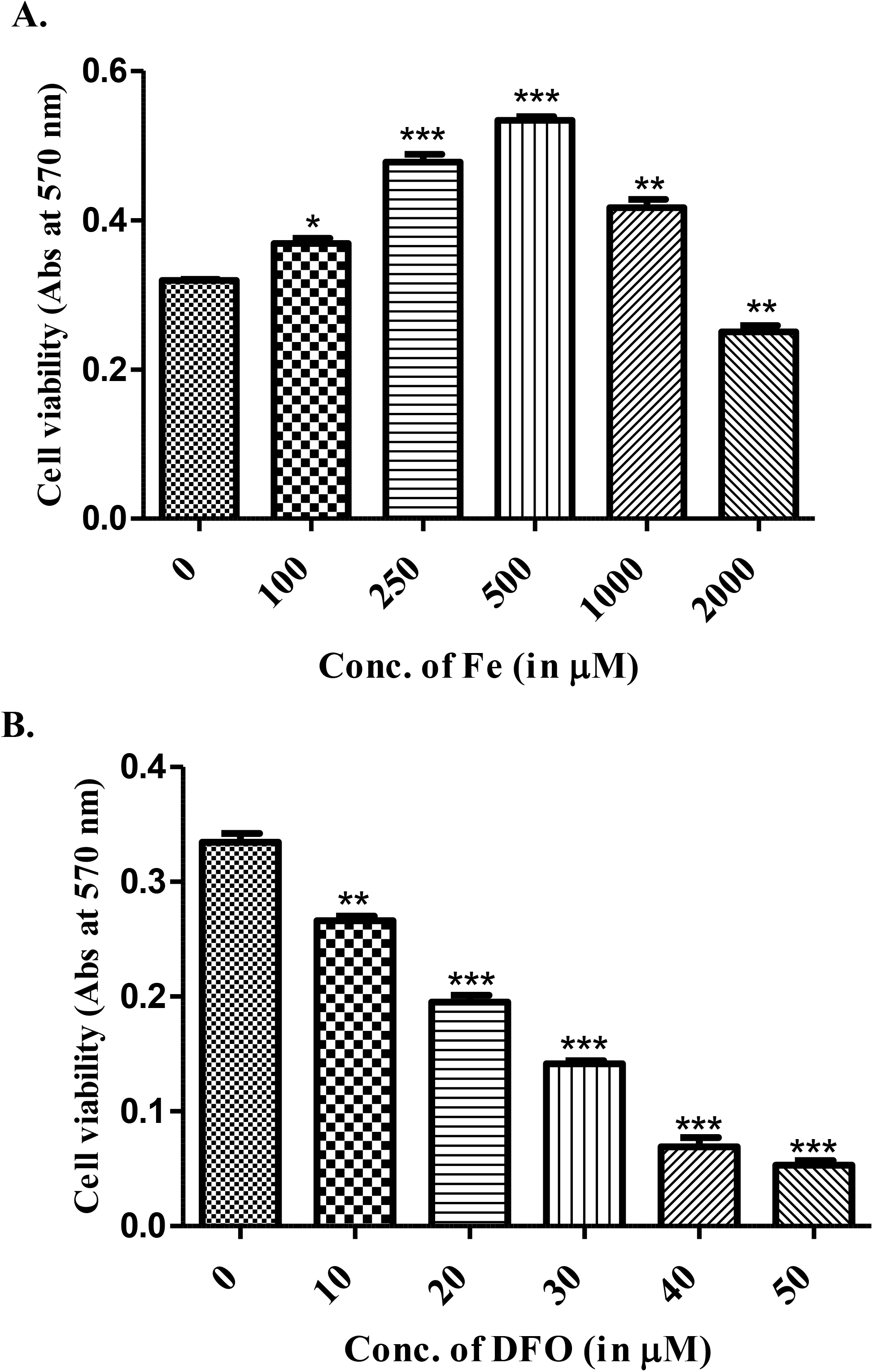
Confirmation of cell viability by MTT assay. Iron Supplemented and depleted promastigote viability was further determined by MTT assay. 2×10^7^ cells/ml were taken and incubated with MTT dye for 4 hrs and absorbance was recorded at 570 nm. The data represents mean ± SEM. Three individual experiments were performed in duplicate. Statistical analysis showed p < 0.001***, p < 0.01** and p < 0.05* performed using one-way ANOVA followed by Tukey’s test.

### 3.6. Iron modulates the infectivity of *L. donovani*

To further validate the impact of Fe-supplementation and depletion upon parasites infectivity or virulence of promastigotes of *L. donovani* were initially treated with Fe_2_S0_4_ꞏ7H_2_O (500 & 1000 μM, Fe-supplemented) **(Fig. 7A)** and DFO (20 & 50 μM Fe-depletion) **(Fig. 8A)**. These parasites were used to infect THP-1-derived macrophages to assess whether iron availability alters parasite infectivity. After 12 hrs of co-incubation (denoted as 0 hrs in Fig 7B & 7C) of both THP-1 macrophages and Fe treated parasites, 500 μM showed higher percentage infectivity (57.4 ± 1 %) with 2.8-fold increase; p < 0.001*** & in 1000 μM Fe showed significantly higher percentage infectivity (92.0 ± 4.8%) with 4.1-fold increase; p < 0.001*** as compared to control group (20.2 ± 1.4%). After 24 hrs of post-infection, 500 μM exhibited higher percentage infectivity (77.8 ± 2.6%) with 3.1-fold increase and 1000 μM Fe parasites exhibited (91.8 ± 2.6%) with increased 3.5-fold change (p < 0.001***) in comparison with control (24.2 ± 2.2%) group. After 48 hrs of post infection, 500 μM Fe showed significantly higher infectivity (72.4 ± 7.2%) with significant fold change of 9.1-fold (p < 0.001***) and 1000 μM Fe parasites exhibited (58.4 ± 5%) higher infectivity with significantly increased 7.3-fold with p value < 0.001*** as compared to untreated control (8.8 ± 1%) **(Fig. 7B)**. After 12 hrs of coincubation (denoted as 0 hrs in Fig 7B & 7C) of macrophages and parasites, the number of amastigotes/100 infected THP-1 macrophage was 362.71 ± 4.41 at 500 μM with increased 3-fold change (p < 0.001***) and 370.80 ± 35.4 at 1000 μM with 3.2-fold increase; with p < 0.001*** in iron supplemented parasites as compared to control 112.12 ± 19.36, respectively. After 24 hrs, no. of amastigotes/100 infected macrophages were 608.57 ± 30.97 for 500 μM with higher 3.9-fold change (p < 0.001***) and 669.06 ± 15.24 for 1000 μM Fe supplemented parasites with increased 4.3-fold; p < 0.001*** in comparison to untreated control 164.46 ± 12.19, respectively. After 48 hrs, no. of amastigotes/100 infected THP-1 macrophages was found 526.79 ± 42.37 for 500 μM iron with 4.4-increased fold change (p < 0.001***) and 661.30 ± 59.78 for 1000 μM iron with 5.5-fold increase (p < 0.001***) in comparison of control 134.09 ± 38.64, respectively **(Fig. 7C)**. The Fe-supplemented parasites showed a substantial increase in the association index and infectivity. These findings indicate a pronounced enhancement in the virulence of the parasites under Fe-supplemented conditions. So, our result clearly indicates that iron availability significantly impacts virulency and leads to better interaction and higher infection of promastigotes of *L. donovani* and host macrophages. To evaluate the role of iron chelation, promastigotes treated with deferoxamine (DFO), an extracellular iron chelator, led to a notable decrease in interaction with THP-1-derived macrophages **(Fig. 8A)** as shown in figures. Both Percent infectivity and parasite load was significantly reduced compared to control.

**Fig 7.**
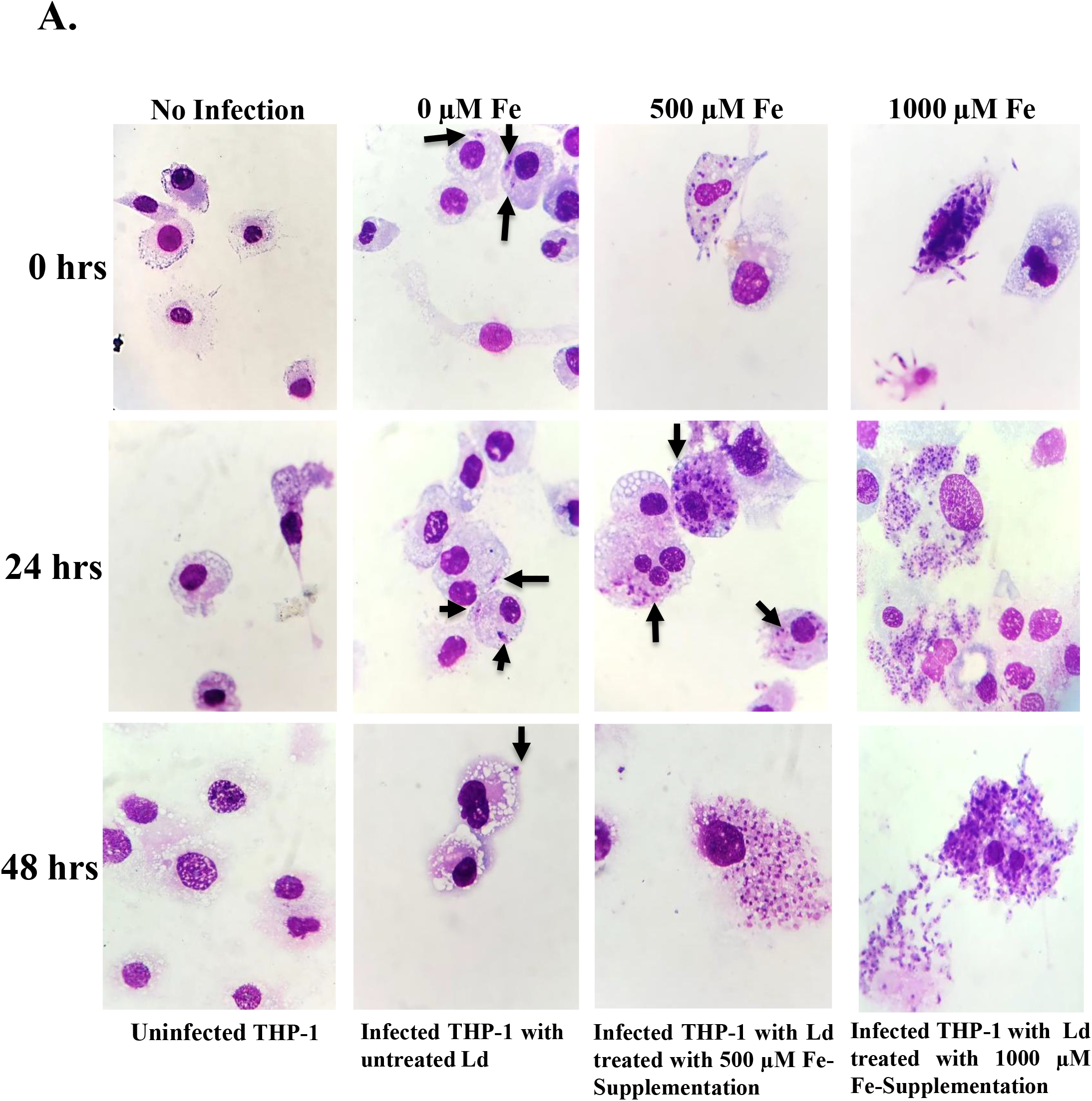
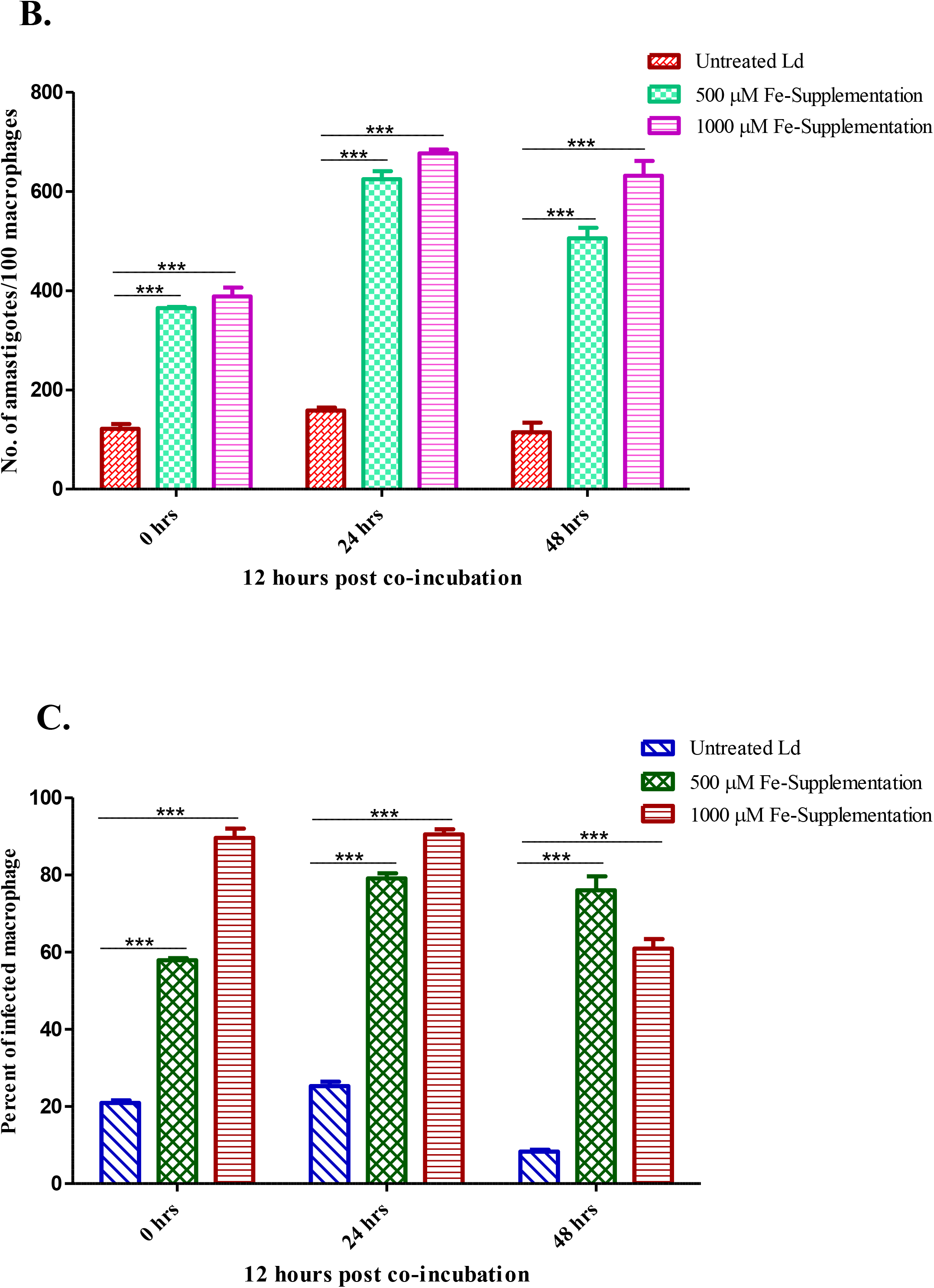
Intracellular iron supplementation induces the interaction between the host macrophage and *L. donovani*. THP-1 derived macrophages were infected with untreated and iron treated *L. donovani* parasites for mention time-period. Parasite was treated with 500 and 1000 µM Fe_2_SO_4_ for Fe-supplementation. **(A)** The image showed THP-1 macrophages infectivity using iron supplemented parasites. **B.** The Rate of infection was determined by counting the no. of amastigotes per 100 macrophages. **C.** The percentage of amastigotes in infected macrophages was determined by counting THP-1 macrophages infected with iron supplemented parasites. The image of infectivity was captured after 12 hrs post-infection and intracellular amastigotes were visualised by Giemsa staining followed by optical microscopy at 100x resolution. The Data represents mean ± SEM of three independent experiments in duplicate. Black arrow indicates intracellular amastigotes in THP-1 macrophages. Statistical analysis showed **p < 0.01 and ***p < 0.001 as compared to untreated (0 µM) parasites by ANOVA as mentioned in materials and methods.

**Fig 8.**
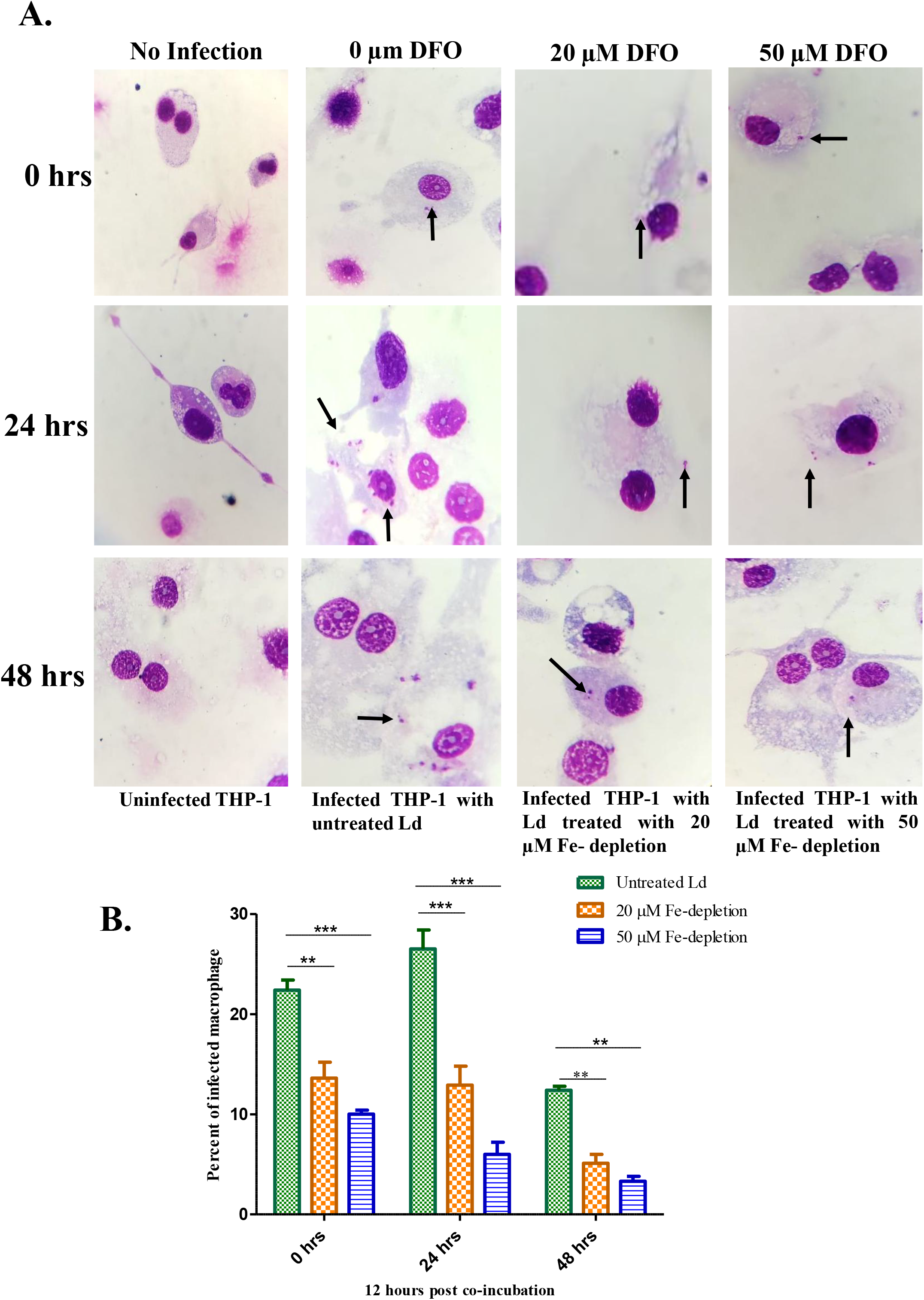
Intracellular iron depletion minimizes the interaction between the host macrophage and *L. donovani*. THP-1 derived macrophages were infected with untreated and DFO treated *L. donovani* parasites for mention time-period. Parasite was treated with 20 and 50 µM DFO for iron depletion. **(A)** The image showed THP-1 macrophages infectivity using iron depleted parasites. **B.** The percentage of amastigotes in infected macrophages was determined by counting infected THP-1 macrophages by iron depleted parasites. The Rate of infection was not determined by counting the macrophages as the no. of amastigotes per 100 macrophages were not determined due to reduced infectivity. The image of infectivity was captured after 12 hrs post-infection and intracellular amastigotes were visualised by Giemsa staining followed by optical microscopy at 100X. The Data represents mean ± SEM of three independent experiments in duplicate. Black arrow indicates intracellular amastigotes in THP-1 macrophages. Statistical analysis showed **p < 0.01 and ***p < 0.001 as compared to untreated (0 µM) parasites by ANOVA test as mentioned thoroughly in material and method section..

After 12 hrs of coincubation, parasites treated with 20 µM DFO demonstrated (12 ± 15.2%) with 1.6-fold decrease (p < 0.01**) in the percentage infectivity in THP-1 macrophages and 50 µM DFO also showed lower percentage infectivity (9.6 ± 0.8%) with 2.21-fold decrease; p value < 0.001(***) in comparison to control (23.4 ± 2%), respectively. After 24 hrs of post infection, parasite treated with 20 µM DFO similarly followed the infection pattern and showed decreased percentage infectivity (11.0 ± 3.8%) with 2.0-fold change (p < 0.001***) and in 50 µM DFO treated parasite exhibited significantly decrease in percentage infectivity as compared to control parasites (24.6 ± 3.8%), respectively. After 48 hours of post infection, result demonstrated decreased percentage infectivity in both 20 µM DFO (6.0 ± 1.8%) with 2.0 fold change decrease (p < 0.001***) and 50 µM DFO simultaneously exhibited reduced percentage infectivity (3.8 ± 1%) with 3.10-fold change; p-value < 0.001(***) as comparison to control group (12.0 ± 3.2%), respectively **(Fig. 8B)**. However, the no. of amastigotes per 100 macrophages was not determined due to lower macrophage infection in 20 and 50 µM DFO. Similarly, parasites treated with 20 and 50 µM DFO showed consistent reductions in the association index of parasite and macrophages and depicted parasite load, indicating an iron-dependent effect on parasite virulence **(Fig 8A)**. In contrast, Fe supplementation significantly enhanced the interaction between THP-1-derived macrophages and promastigotes of *L. donovani*. The decrease in parasite load and infectivity observed with DFO-treated parasites underscores the crucial role of iron in the pathogenicity of *L. donovani*. DFO, by chelating extracellular iron, impairs the ability of the parasites to interact with and infect macrophages.

Our result highlights the essential role of iron in maintaining parasite virulence and infection of macrophages. The significant enhancement in parasite load and percentage infectivity in Fe-supplemented conditions suggests that iron availability can augment the virulence of promastigotes of *L. donovani* and further supports the notion that iron is a critical factor for parasite proliferation within macrophages. Overall, our work emphasizes that iron is crucial factor in *L. donovani* infection: Fe-depletion reduces infectivity and virulence, while Fe-supplementation enhances parasite macrophage interaction and intracellular amastigotes proliferation.

### 3.7. Iron supplementation and depletion promotes apoptosis mediated cell death in *L. donovani*

*L. donovani* parasite shows a tightly regulated iron acquisition and transportation mechanism [39]. Our experimental data revealed that external iron treatment disturbs the iron homeostasis and cellular redox balance. So, we verified whether an excess or low iron condition induced LD cell death. The results from Fe supplementation showed a prominent increase in caspase-like activity in a concentration-dependent manner. We estimated caspase-like activity in the presence of iron, where we found a ̴ 1.1-fold, ̴ 1.6-fold, ̴ 2-fold increase at 500, 1000 & 2000 µM which revealed higher iron concentration leads to increased intracellular stress **(Fig. 9A)**. Iron depletion also led to increased caspase-like activity in concentration concentration-dependent manner, where we found a ̴ 1.4-fold, ̴ 1.9-fold, ̴ 2.1-fold increase at DFO conc. of 10 µM, 30 µM, & 50 µM **(Fig. 9B)**. These findings suggest that iron can also induce apoptosis-like events in promastigotes of *L. donovani*. DNA fragmentation was used to further analyze the hallmarks and decisive factors of apoptosis, which includes the activation of proteases DNA fragmentation was used to further analyze the hallmarks and decisive factors of apoptosis, which include the activation of proteases and evidence of genomic DNA degradation. Therefore, we isolated DNA from both iron-supplemented and depleted cells and performed agarose gel electrophoresis to visualise DNA fragmentation. We detected that the higher concentrations of Fe_2_SO_4_ (1000 µM and 2000 µM) led to DNA fragmentation **(Fig. 9C),** while treatment with DFO 10, 20, 30, 40, and 50 µM led to DNA fragmentation in cells **(Fig. 9D),** further suggesting that excess or low iron led to apoptosis.

**Fig 9.**
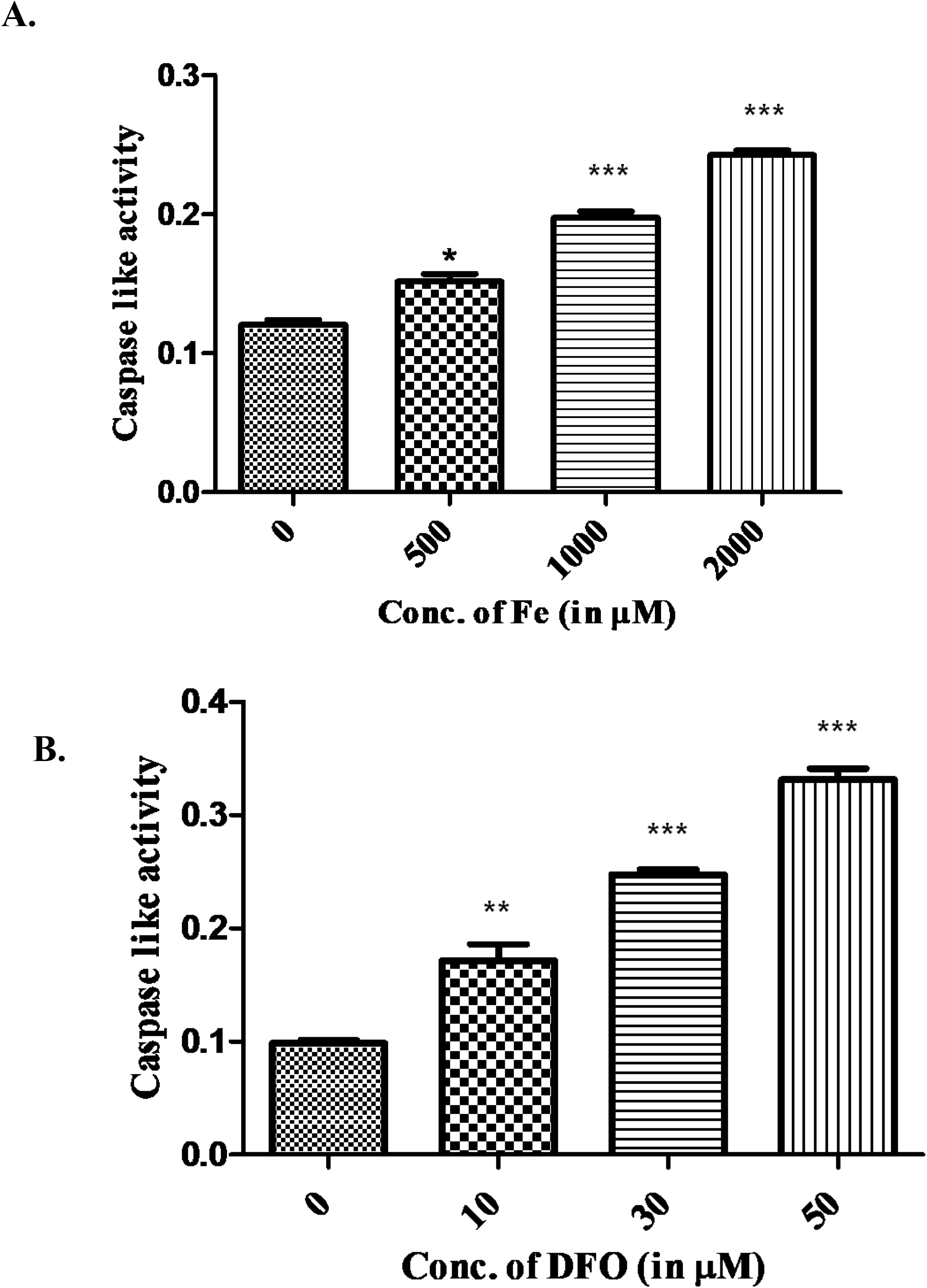
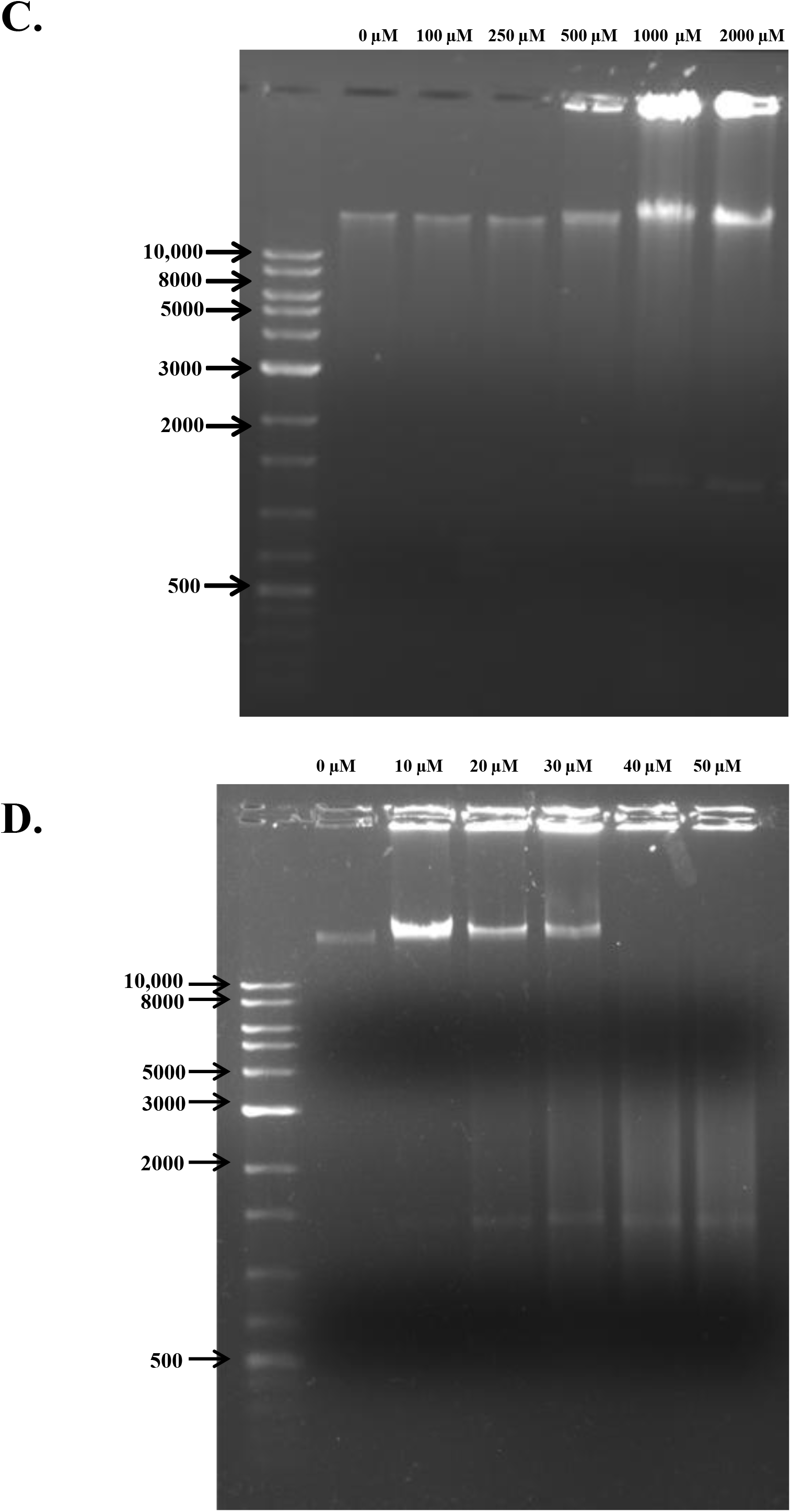
Excess or low intracellular iron level leads to the cell death. Caspase-like activity was determined in **A**. iron supplemented (500, 100, and 2000 µM) and **B.** iron depleted (10, 30, and 50 µM) parasites of *L. donovani* as mentioned in method and material section. The triplicate individual experiment was carried out. Data represent mean ± SEM of three independent experiment, statistically significant values are denoted as p < 0.05*, p < 0.01**, p < 0.001*** by t-test when compared with untreated. Manual DNA isolation was performed for analysis of DNA fragmentation in **C.** iron supplemented and **D.** iron depleted condition. Lane descriptions are mentioned above the gel.

### 3.8. Intracellular iron alters parasite redox balance

ROS are typically generated as a by-product of disrupted iron metabolism and can cause cellular damage if not properly managed [40, 41]. To further validate the involvement of excess or low iron associated with oxidative stress in the WT strain, we estimated ROS and total intracellular thiol. The intracellular ROS content in the WT parasites was measured using H_2_DCFDA dye. The amount of fluorescence generated is closely related to intracellular reactive oxygen species. In iron supplementation condition, ROS was found decreased ̴ 1.1-fold; p < 0.01** at 100 µM, ̴ 1.2-fold; p < 0.01** at 250 µM, ̴ 1.2-fold; p < 0.001*** at 500 µM, ̴ 1.4-fold; p < 0.001*** at 1000 µM Fe conc., ̴ 1.5-fold; p < 0.001*** at 2000 µM Fe conc. and ̴ 1.9 fold (p < 0.001***) at 4000 µM **(Fig. 10A)**. Our findings suggest that reduced ROS production as a result of intracellular Fe supplementation may be a source of induced viability in Fe-supplemented parasites. However, we have observed a gradual increase in intracellular total thiol content suggesting ̴ 1.5-fold; p < 0.01** at 100 µM, ̴ 2-fold; p < 0.001*** at 250 µM, ̴ 2.6-fold; p < 0.001*** at 500 µM, ̴ 3-fold; p < 0.001*** at 1000 µM, ̴ 3.6-fold; p < 0.001*** at 2000 µM and ̴ 4.4-fold; p < 0.001*** at 4000 µM in Fe-supplemented promastigote parasites **(Fig. 10B)**.

**Fig 10.**
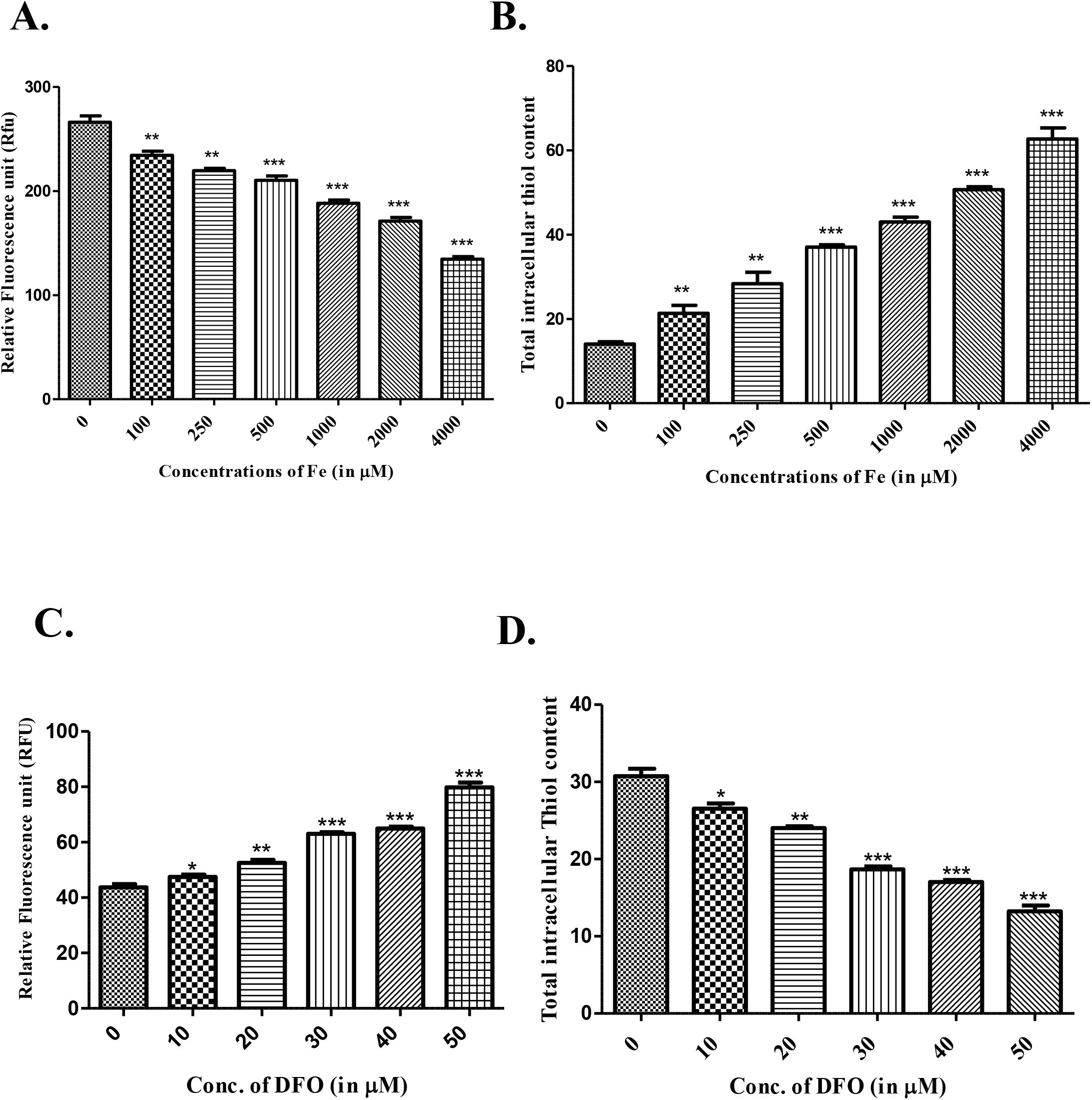
Iron affects ROS and total intracellular thiols in the parasites. Treated and untreated control parasites were cultured in M199 medium for 5 days. Intracellular accumulation of ROS was estimated using 2, 7-dichlorodihydrofluorescein diacetate (H_2_DCFDA) by using a spectrofluorometer and absorbance was recorded at an excitation wavelength of 504 nm and emission of 529 nm and denoted as Relative fluorescence unit (RFU). The amount of RFU is directly proportional to the recorded fluorescence intensity. The total intracellular reduced thiol content was estimated using DTNB and absorbance measured at 412 nm by using a spectrophotometer and denoted as micromoles (µ-moles). **A & B.** ROS and total intracellular thiol content WT strain cultured in iron supplemented conditions from 0 µM to 4000 µM. **C & D.** ROS and total intracellular thiol content was estimated in WTAg83 strain of *L. donovani* cultured in iron-depleted conditions from 0 µM to 50 µM. The individual triplicate experiment was performed, and data represent mean ± SEM *p < 0.05, **p < 0.01, ***p < 0.001 statistically analysed by using Students t-test.

Our results suggest changes in total thiol content in parasites are 2 to 3-fold higher than the decrease in ROS level decreases in the parasites. The Fe-supplemented parasites have higher intracellular thiol levels, which are correlated with lower ROS levels and higher parasite survival. We also assessed how iron deficiency affected the production of oxidative stress in parasites. Our result showed ̴ 1.1-fold; p < 0.05*, ̴ 1.2-fold; p < 0.01**, ̴ 1.6-fold; p < 0.001***, ̴ 1.7-fold; p < 0.001***, ̴ 2.1-fold; p < 0.001*** decrease in total intracellular thiol content at 10, 20, 30, 40 & 50 µM Fe-depleted parasites **(Fig. 10D)**. ROS estimation in iron depleted condition showed ̴ 1.1-fold with p < 0.05*, ̴ 1.2-fold with p < 0.01**, ̴ 1.3-fold with p < 0.001***, ̴ 1.5-fold with p < 0.001*** & ̴ 1.52-fold with p < 0.001*** increase at conc. 10, 20, 30, 40 & 50 µM **(Fig. 10C)** depicts that iron depletion induced reactive oxygen species (ROS) in WT Ag83 strain provides important insights into the oxidative stress responses of *Leishmania* parasites under iron deficient conditions. The induction of ROS upon iron depletion is indicative of the parasite’s response to reduced iron availability, which disrupts normal cellular iron-dependent processes and leads to ROS which is required for the proliferation of the cells upto certain concentration. Since, cells proliferation gets hampered due to low iron availability, so parasites try to cope up with this low iron situation and further required support of intracellular total thiol level but production of total reduced intracellular thiol is found low in iron depleted conditions. Thus, Fe-depleted parasites may not be able to scavenge increased ROS formation due to reduced intracellular thiol, which dropped the parasite’s chances of survival. A comparison in ROS levels and total intracellular thiol content observed under excess or low iron conditions clearly emphasizes a strong positive correlation among them, which is aptly supported by the fact that ROS can generate oxidative damage, and a lower thiol level is not sufficient to scavenge ROS in depleted conditions, leading to cell death.

### 3.9. Iron supplementation mediated tolerance toward oxidative stress

By subjecting *Leishmania* parasites to menadione-induced oxidative stress in both iron-supplemented and iron-depleted environments, we examined the function of iron in oxidative stress tolerance. WT promastigotes were incubated for 0-18 hrs, and at 3-hr time point, the parasite’s percentage cell viability was calculated. Our findings demonstrated a striking difference in oxidative stress tolerance depending on iron availability. Iron-depleted parasites (DFO-treated) showed significantly reduced survival upon exposure to menadione, failing to tolerate even the lower concentration of menadione (2.5 µM). However, iron-supplemented parasites exhibited better tolerance to oxidative stress, surviving both 2.5 µM **(Fig. 11A)** and 5.0 µM **(Fig. 11B)** menadione concentration. However, our findings suggest the different impact of both 2.5 and 5.0 µM menadione concentration on iron supplemented parasites (0-4000 µM) and we observed clear difference in the percentage viability were 5.0 µM menadione concentration shows higher growth inhibition.

**Fig 11.**
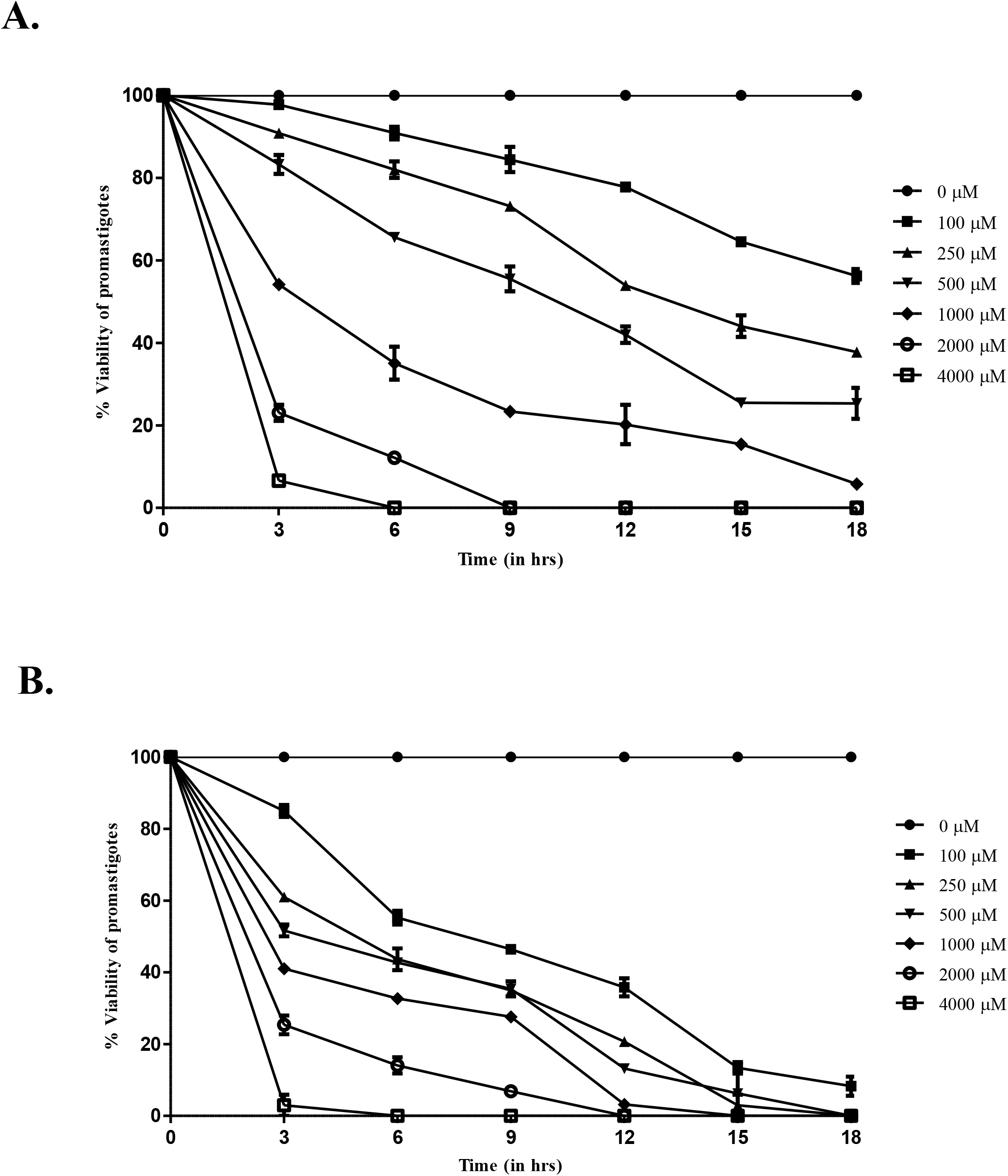
Iron supplementation increases tolerance toward oxidative stress. Percentage viability curve of WT promastigotes cultured in iron supplemented condition after exposed to (**A)** 2.5 µM and (**B)** 5.0 µM concentration of menadione. The percentage viability was determined after every 3 hrs of time interval for 18-hrs incubation period. Data represented mean ± SEM values. Three individual experiments were performed in triplicate and analysed using Student’s t-test.

At 3 hrs, the parasites treated with 2.5 µM menadione showed 98.56 ± 1.56% cell viability at 100 µM Fe, 90.67 ± 1.67% viability at 250 µM Fe, 85.56 ± 4.56% viability at 500 µM Fe, 53.33 ± 1.67% viability at 1000 µM Fe, 21.11 ± 3.89% viability at 2000 µM Fe and significantly reduced 5.82 ± 1.57% viability showed more tolerance toward oxidative stress in comparison to control. After 6 hrs with 2.5 µM menadione, percentage viability of parasites reduced to 92.44 ± 3.08% cell viability at 100 µM Fe, 84.0 ± 4% viability at 250 µM Fe, 66.67 ± 2.12% viability at 500 µM Fe, 31.11 ± 7.95% viability at 1000 µM Fe, 11.0 ± 2.30% viability at 2000 µM Fe and at 4000 µM Fe, the percentage viability decreased to 0%, respectively. After 9 hrs, in presence of 2.5 µM menadione; the parasite cultured in 2000 & 4000 µM Fe at higher iron supplementation condition couldn’t survive and growth was inhibited. Iron supplemented parasites from 100-1000 µM survived oxidative stress induced by 2.5 µM menadione and managed to survive managed to survive 18 hrs. After 18 hrs, the percentage viability was 57.89 ± 3.34% at 100 µM, 37.37 ± 0.83% at 250 µM, 21.58 ± 7.52% at 500 µM and significantly reduced to 5.79 ± 0.03% at 1000 µM in comparison to control **(Fig. 11A)**. The Fe-Supplemented parasites treated with 5.0 µM menadione showed significantly lower oxidative stress tolerance and increased growth inhibition at higher iron concentration. At 3 hrs, parasite treated with 5.0 µM menadione exhibited 83.33 ± 3.34% viability at 100 µM, 60.0 ± 2% viability at 250 µM, 53.33 ± 3.0% viability at 500 µM, 40.0 ± 2.09% viability at 1000 µM, 22.76 ± 5.2% viability at 2000 µM and significantly low 5.88 ± 0.0% viability at 4000 µM as compared to control. After 6 hrs, parasite treated with 5.0 µM menadione showed 57.14 ± 3.81% viability at 100 µM, 40.63 ± 6.04% viability at 250 µM, 41.43 ± 2.57% viability at 500 µM, 34.0 ± 2.6% viability at 1000 µM, 11.82 ± 4.51% viability at 2000 µM as compared to control and for 4000 µM, higher cytotoxicity led to cell death after 6 hrs in presence of 5.0 µM menadione. After 9 hrs, the percentage viability reduced to 46.0 ± 0.88% for 100 µM, 36.0 ± 2% for 250 µM, 37.50 ± 4.17% for 500 µM, 28.57 ± 1.9% for 1000 µM, 5.88 ± 1.94% for 2000 µM and no viability found at 4000 µM in comparison to control. After 18 hrs, lower iron concentration 100 µM parasites managed to survived stress which showed significantly reduced 10.95 ± 5.34% percentage viability as compared to control **(Fig. 11B)**. The menadione oxidative stress result revealed remaining ̴ 80% cell viability at 2.5 µM menadione & ̴ 48% cell viability at 5.0 µM menadione at 9 hrs in 100 µM Fe, ̴ 70% cell viability at 2.5 µM menadione & ̴ 35% cell viability at 5.0 µM menadione at 9 hrs in 250 µM Fe, ̴ 55% cell viability at 2.5 µM menadione & ̴ 35% cell viability at 5.0 µM menadione at 9 hrs in 500 µM Fe, ̴ 23% cell viability at 2.5 µM menadione & ̴ 26% cell viability at 5.0 µM menadione at 9 hrs in 1000 µM Fe, Parasite growth inhibited at 2.5 µM menadione at 9 hrs in 2000 µM Fe & showed significantly reduced ̴ 6% cell viability at 5.0 µM menadione in compared to control. While at 12 hrs, result exhibited remaining ̴ 77% cell viability at 2.5 µM menadione, ̴ 35% cell viability at presence of 5.0 µM menadione in 100 µM Fe, ̴ 53% cell viability at 2.5 µM menadione & ̴ 20% cell viability at 5.0 µM menadione in 250 µM Fe, ̴ 42% cell viability at 2.5 µM menadione & ̴ 12% cell viability at 5.0 µM menadione in 500 µM Fe, ̴ 18% cell viability at 2.5 µM menadione & significantly lower ̴ 3% cell viability at 5.0 µM menadione at 1000 µM Fe, meanwhile no cell viability at 2000 & 4000 µM Fe in both 2.5 & 5.0 µM menadione as compared to control showed complete growth inhibition, respectively. Further, at 15 hrs of menadione conc. comparison data showed a drastic difference where the cell viability at 2.5 µM menadione was ̴ 64% & at 5.0 µM menadione the cell viability reduced to ̴ 13% at 100 µM Fe-supplementation, ̴ 43% cell viability at 2.5 µM menadione & ̴ 5% cell viability at 5.0 µM menadione in 250 µM Fe-supplementation which also represented a huge difference, ̴ 25% cell viability at 2.5 µM menadione & ̴ 12% cell viability at 5.0 µM menadione in 500 µM Fe-supplementation, ̴ 20% cell viability at 2.5 µM menadione & no parasite viability was found at 5.0 µM menadione in 1000 µM Fe in comparison to control, respectively **(Fig. 11A & B)**. Our result showed cell viability was decreasing with increasing concentration of menadione and iron supplementation. Menadione and iron supplementation were both found to be associated with percentage cell viability, and higher menadione with elevated concentration of iron led to increased intracellular stress and cytotoxicity. Our data suggested that iron supplementation has a critical role in oxidative stress management in *L. donovani* promastigotes, likely by supporting redox balancing mechanisms essential for parasite survival under hostile conditions.

### 3.10. Iron also modulates Fe-S and thiols proteins expression

To validate the impact of intracellular iron after Fe supplementation (0-4000 µM Fe_2_SO_4_) and depletion (0-50 µM DFO) on *L. donovani* promastigotes, we compared the expression level of protein involved in thiol metabolic pathway and Fe-S cluster assembly proteins. Our result demonstrated the expression of LdTryS, LdTryR, LdCS, LdSAT, LdcTXN, LdcTXNPx, LdFrataxin and LdIscU at protein level was significantly higher in iron supplemented condition than in wild-type control parasites except of LdNbp35 protein. The careful densitometric analysis revealed the fold change difference at expression level of protein on different iron conditions as compared to control protein (LdActin). In iron supplemented condition for LdTryS, we observed increased ̴ 1.2-fold change at 100 & 250 µM, increased ̴ 1.3-fold change at 500 & 1000 µM, increased ̴ 1.5-fold change at 2000 µM and increased ̴ 1.6-fold change at 4000 µM. Our data revealed that LdTryS protein expression was elevated with increasing concentration of iron supplementation and higher LdTryS expression was observed at 4000 µM in comparison to other iron concentration and as compared with wildtype control. For LdTryR, expression revealed increased 1.2-fold at 100 & 250 µM, ̴ 1.4-fold at 500 µM, ̴ 1.5-fold at 1000 µM, ̴ 1.6-fold at 2000 µM and significantly increased ̴ 2.4-fold at 4000 µM with compared to control. Our result showed LdTryR protein expression was affirmatively upregulated in higher iron concentration (4000 µM) as per wildtype control. There were non-significant fold changes at both 100 and 250 µM for LdSAT protein expression. Apart from 100 & 250 µM, other iron concentration showed increase in fold change in expression of LdSAT protein. We found ̴ 1.4-fold increase at 500 µM, ̴ 1.6-fold increase at 1000 & 2000 µM and significantly increased ̴ 2.1-fold at 4000 µM as compared to control shows expression of LdSAT protein also gets upregulated in 4000 µM; higher iron concentration, respectively. Similarly, our result also revealed there was no such difference in protein expression level of LdCS and observed non-significant fold change at initial 100 & 250 µM iron supplementation. The expression level of

LdCS was increased ̴ 1.1-fold at 500 µM, ̴ 1.3-fold at 1000 µM, ̴ 1.5-fold at 2000 µM and ̴ 1.4-fold at 4000 µM as compared to control. For LdNbp35 protein expression, we found ̴ 1.1-fold increase at 100 µM, ̴ 1.4-fold increase at 250 µM, ̴ 1.5-fold increase at 500 µM, ̴ 1.6-fold increase at 1000 µM, ̴ 1.1-fold decrease at 2000 µM and no difference at 4000 µM as compared to control. However, LdFrataxin protein level at lower iron supplementation showed no such difference at 100 and 250 µM, while at 500 µM ̴ 1.1-fold increase, at 1000 µM ̴ 1.2-fold increase, at 2000 & 4000 µM, ̴ 2.1-fold increase as compared to control was found which suggest higher iron supplementation condition upregulates LdFrataxin expression (mitochondrial iron storage protein). The Expression level of LdcTXNPx showed no change at 100 µM, ̴ 1.2-fold increase at 250 & 500 µM, ̴ 1.3-fold increase at 1000 & 2000 µM and ̴ 1.4-fold increase at 4000 µM in comparison to untreated control. There was no fold change difference in expression of LdIScU at initial iron supplementation concentration (100 µM). After this, we observed ̴ 1.1-fold increase at 250 µM, ̴ 1.3-fold increase at 500 µM, ̴ 1.4-fold increase at 1000 µM, ̴ 1.6-fold increase at 2000 µM and ̴ 1.9-fold increase at 4000 µM in the expression level of LdIScU as compared to control. For LdcTXN, the protein expression was ̴ 1.1-fold higher at 100 µM, ̴ 1.4-fold higher at 250 µM, ̴ 1.6-fold higher at 500 µM and ̴ 1.7-fold higher at 1000, 2000 & 4000 µM. The expression pattern of LdcTXN in iron supplementation was initially upregulated and expression level remains constant at higher iron concentration **(Fig. 12A)**. The increased expression of thiol metabolic proteins is associated with the intracellular thiol level and support that total thiol production during oxidative stress for the survival of the parasites is increased.

**Fig 12.**
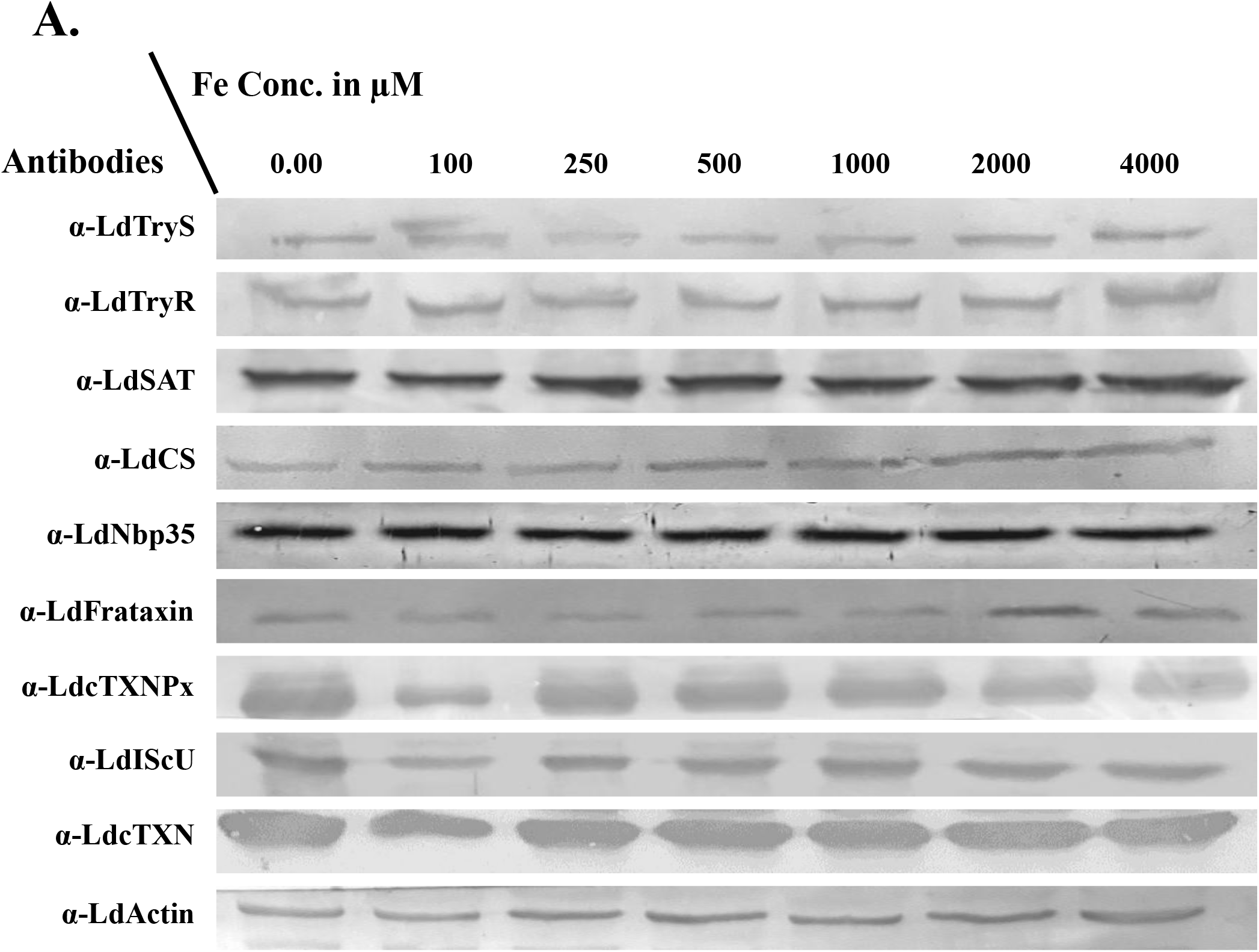
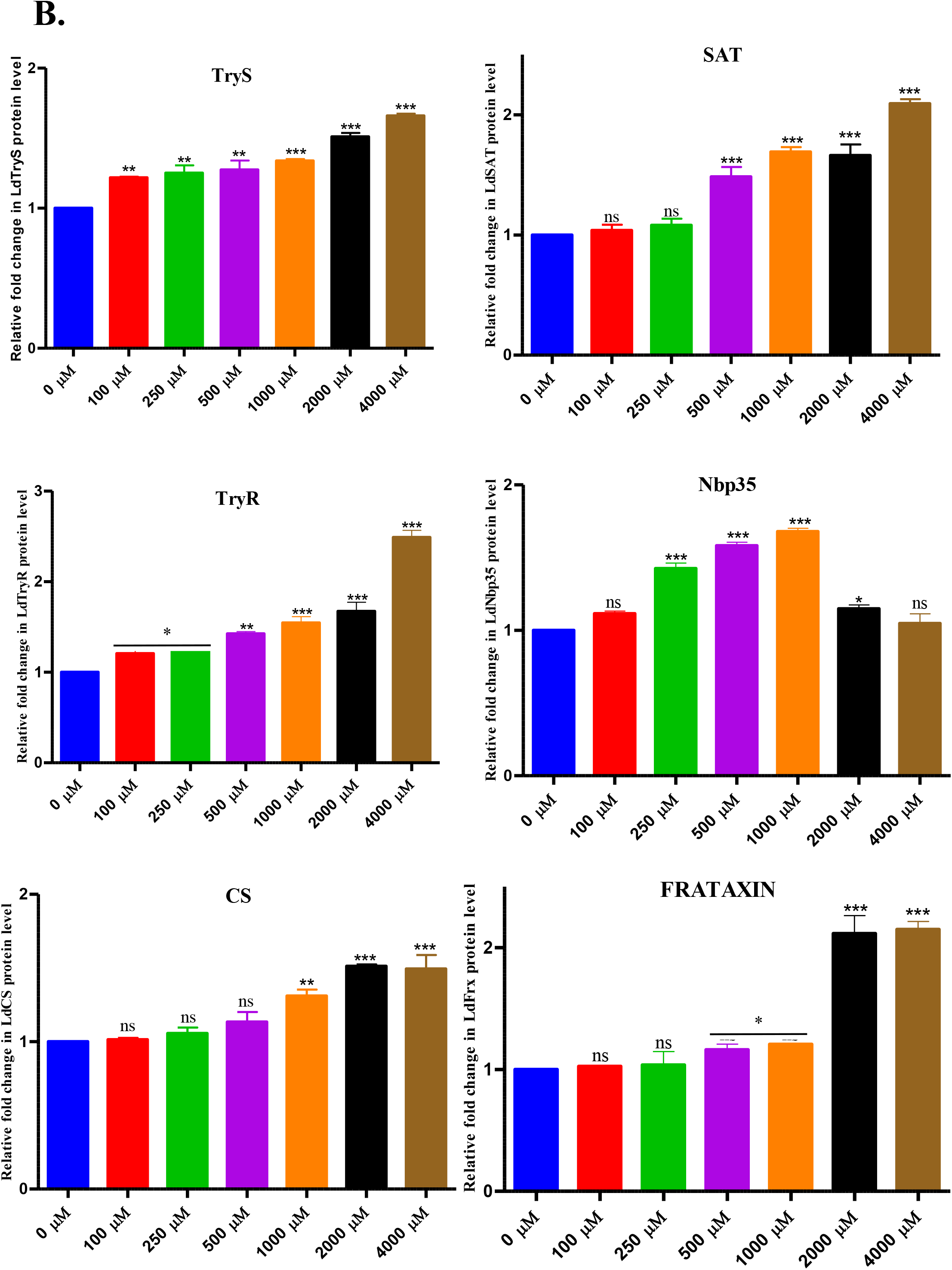
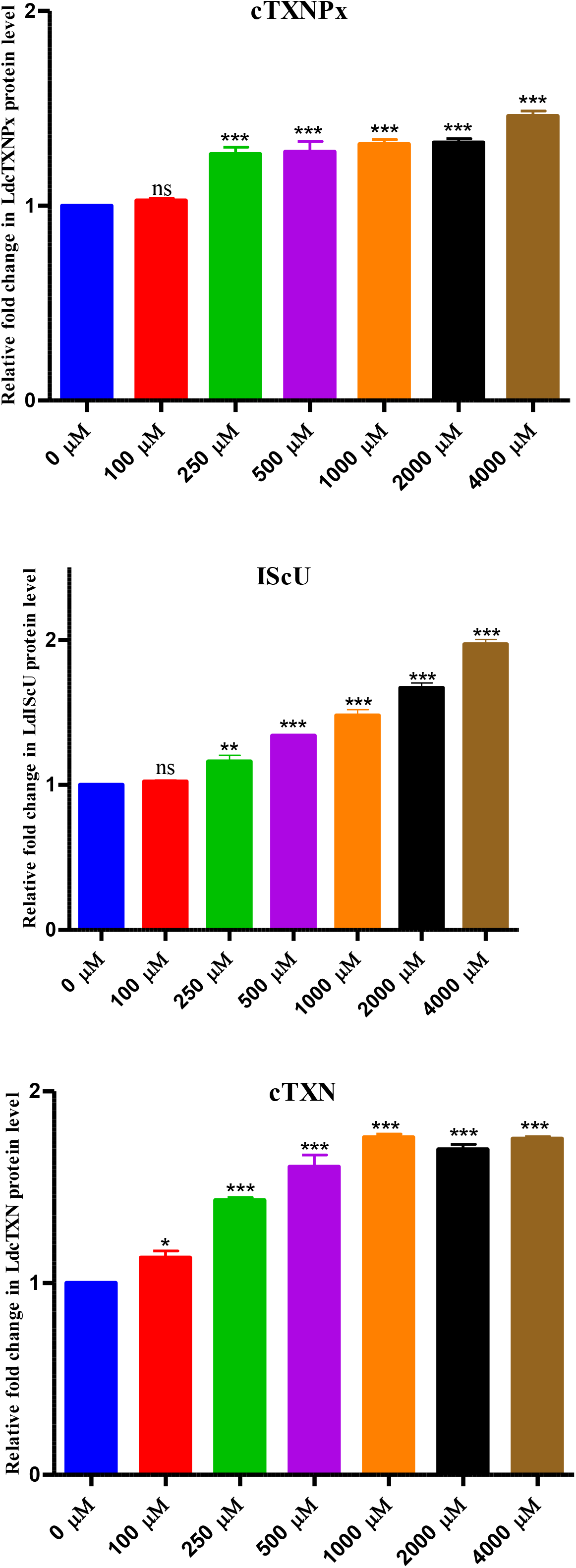
Expression of thiol metabolic and Fe-S cluster proteins analysis and relative fold changes of iron supplemented promastigote parasites. WT promastigotes were treated with different concentrations of iron-supplemented (0-4000 µM) for 5 days. After treatment, 30 µg of total cell lysate was loaded on SDS-gel and blotted onto PVDF membrane and further probed with the mentioned *L. donovani* polyclonal antibodies. For loading control, anti-LdActin was used. The western blotting shows expression of *L. donovani* mentioned proteins involved in thiol metabolism and Fe-S cluster biogenesis in iron-supplemented conditions. Relative band intensities were determined by performing densitometric analysis using ImageJ software and fold change was calculated by normalising the band intensity with loading control. The abundance of thiol and Fe-S proteins in iron-supplemented represented by a graphical plot. Statistical analysis was performed by Student’s t-test and denoted ns for non-significant, *p < 0.05, **p < 0.01, ***p < 0.001 compared to untreated control.

In iron depleted condition (DFO), the densitometric analysis revealed that the majority of thiol and Fe-S proteins expression level significantly downregulated except LdFrataxin. The LdTryS protein was ̴ 1.2-fold decreased at 10 µM, ̴ 1.7-fold decreased at 20 µM, ̴ 3.5-fold decreased at 30 µM, ̴ 4.5-fold decreased at 40 µM and a significant ̴ 6.1-fold decreased at 50 µM as compared to control. The expression level of LdTryR was ̴ 1.2-fold decreased at 10 µM, ̴ 1.3-fold and 1.5-fold decreased at 30 & 40 µM, respectively and ̴ 1.6-fold decreased at 50 µM as compared to control. The LdCS protein expression was ̴ 1.2-fold decreased at 10 µM, ̴ 1.3-fold decreased at 20 µM, ̴ 1.4-fold decreased at 30 µM, ̴ 1.5-fold decreased at 40 µM, and ̴ 1.6-fold decreased at 50 µM. Our result showed the expression level of LdSAT was not changed at 10 & 20 µM DFO treated cells, but expression level was downregulated at 30 µM with ̴ 1.2-fold decrease, 40 & 50 µM with ̴ 1.3-fold decrease, respectively. The protein LdNpb35 expression level was significantly decreased and showed ̴ 1.8-fold decrease at 10 µM, ̴ 2.0-fold decrease at 20 µM, ̴ 2.3-fold decrease at 30 µM, ̴ 2.4-fold decrease at 40 µM and ̴ 3.8-fold decrease at 50 µM as compared to control. However, LdFrataxin protein was upregulated with increasing DFO concentration and showed ̴ 1.1-fold higher expression at 10 µM, ̴ 1.3-fold higher expression at 20 µM, ̴ 1.4-fold higher expression at 30 µM, ̴ 1.6-fold higher expression at 40 µM and ̴ 1.9-fold higher LdFrataxin protein expression at 50 µM as compared to control. Possibly, the upregulation of LdFrataxin was associated with parasite survival and in iron depleted condition in parasites because it uses Frataxin protein as iron reserve to fulfil intracellular iron needs and possibly low intracellular iron condition triggers Fe-S protein pathway and upregulation of the Frataxin protein for the fulfilment of cellular events during iron scarcity. The expression level of LdcTXNPx was also decreased and showed ̴ 1.4-fold lower expression at 10 µM, ̴ 1.7-fold lower at 20 µM, ̴ 2.0-fold lower at 30 µM, ̴ 2.3-fold lower at 40 µM and ̴ 2.2-fold lower protein expression at 50 µM in comparison to control. The expression level of LdIScU protein showed no change at 10 µM, ̴ 1.2-fold decrease at 20 µM, ̴ 1.5-fold decrease at 30 µM, ̴ 1.6-fold decrease at 40 µM and ̴ 1.7-fold decrease at 50 µM as compared to control. Similarly, the LdcTXN protein expression showed no fold change in expression level at 10 µM, ̴ 1.1-fold decrease at 20 & 30 µM, ̴ 1.2-fold decrease at 40 µM, and ̴ 1.7-fold decrease in LdcTXN expression at 50 µM in comparison to control **(Fig. 13A)**.

**Fig 13.**
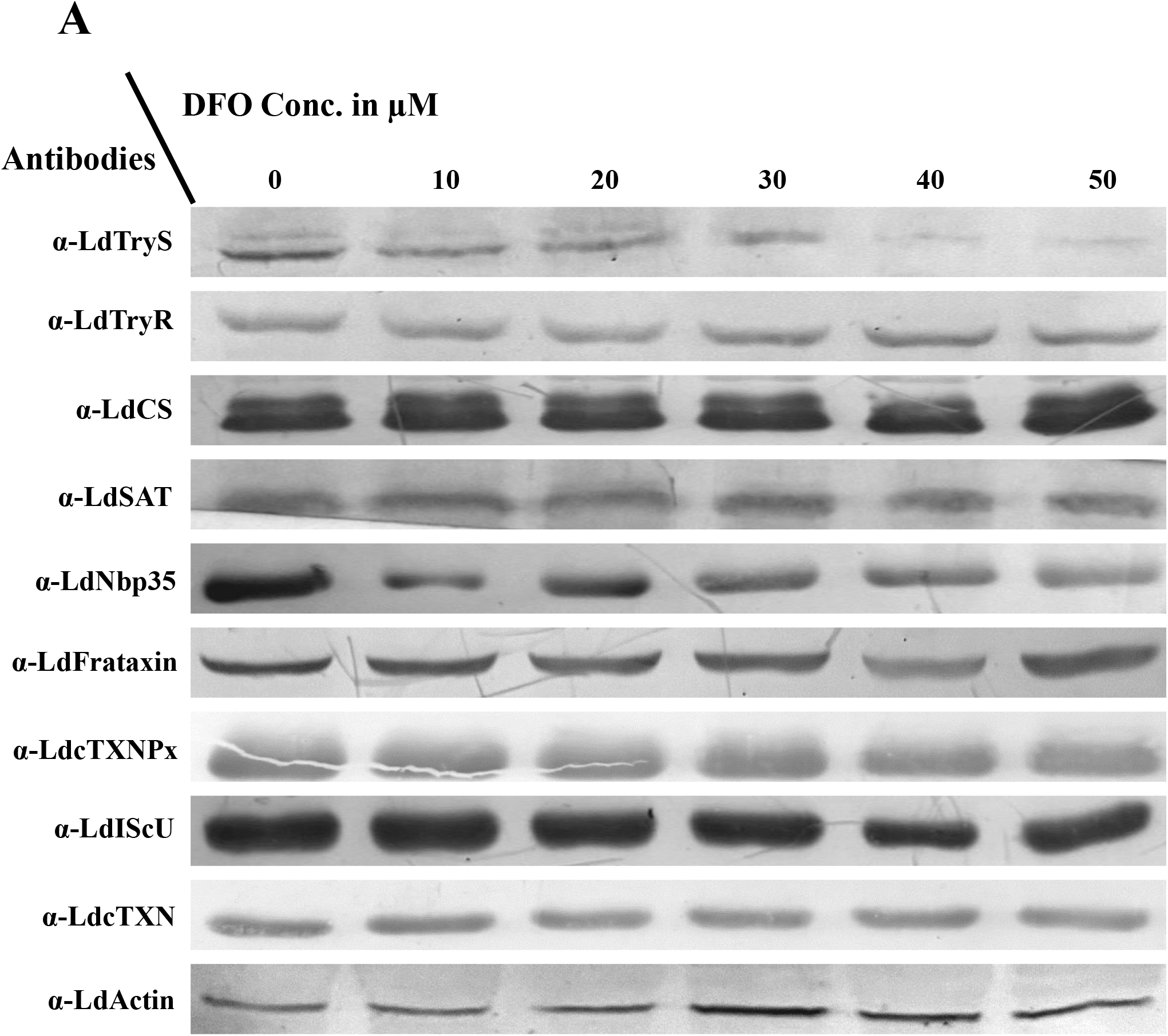
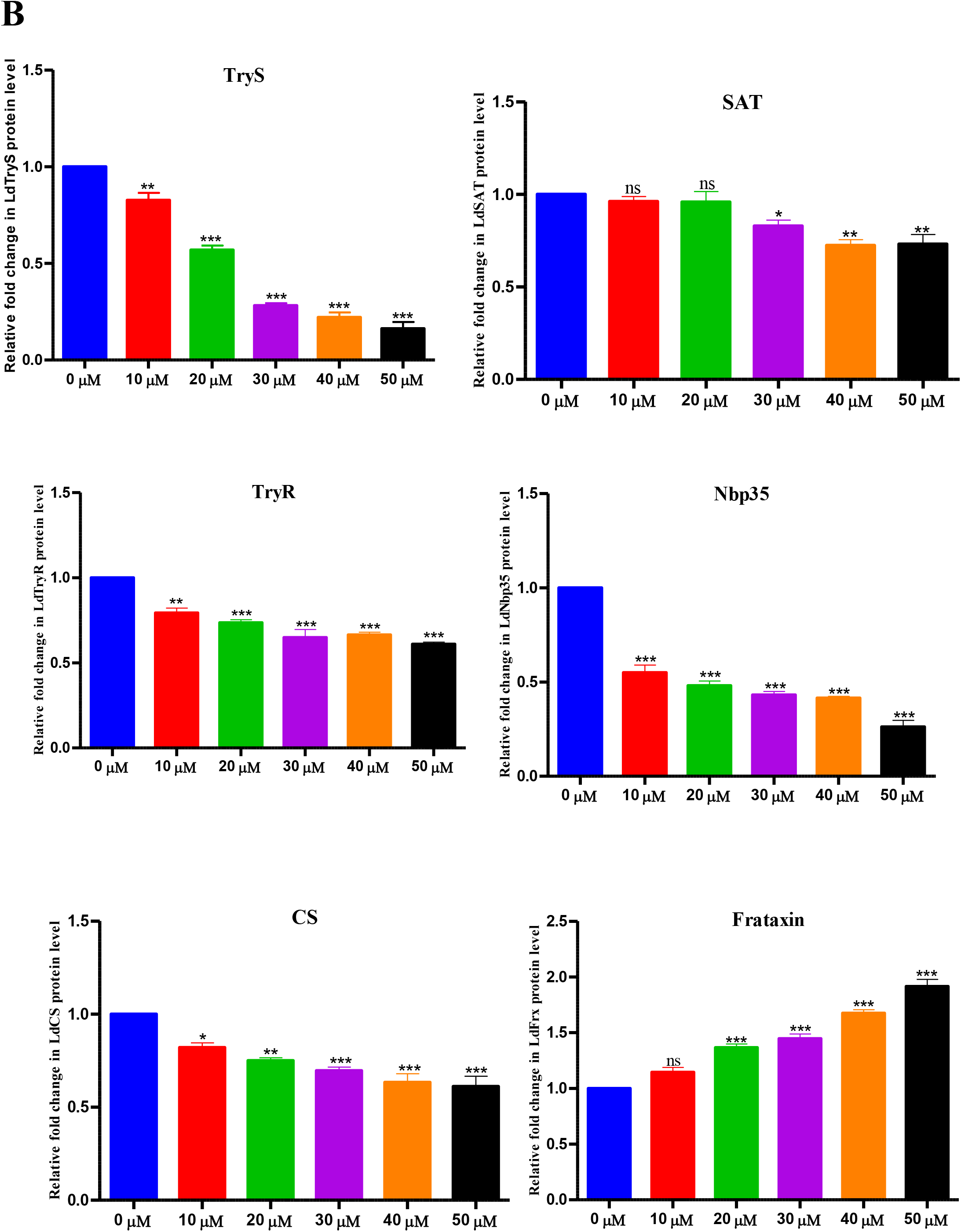
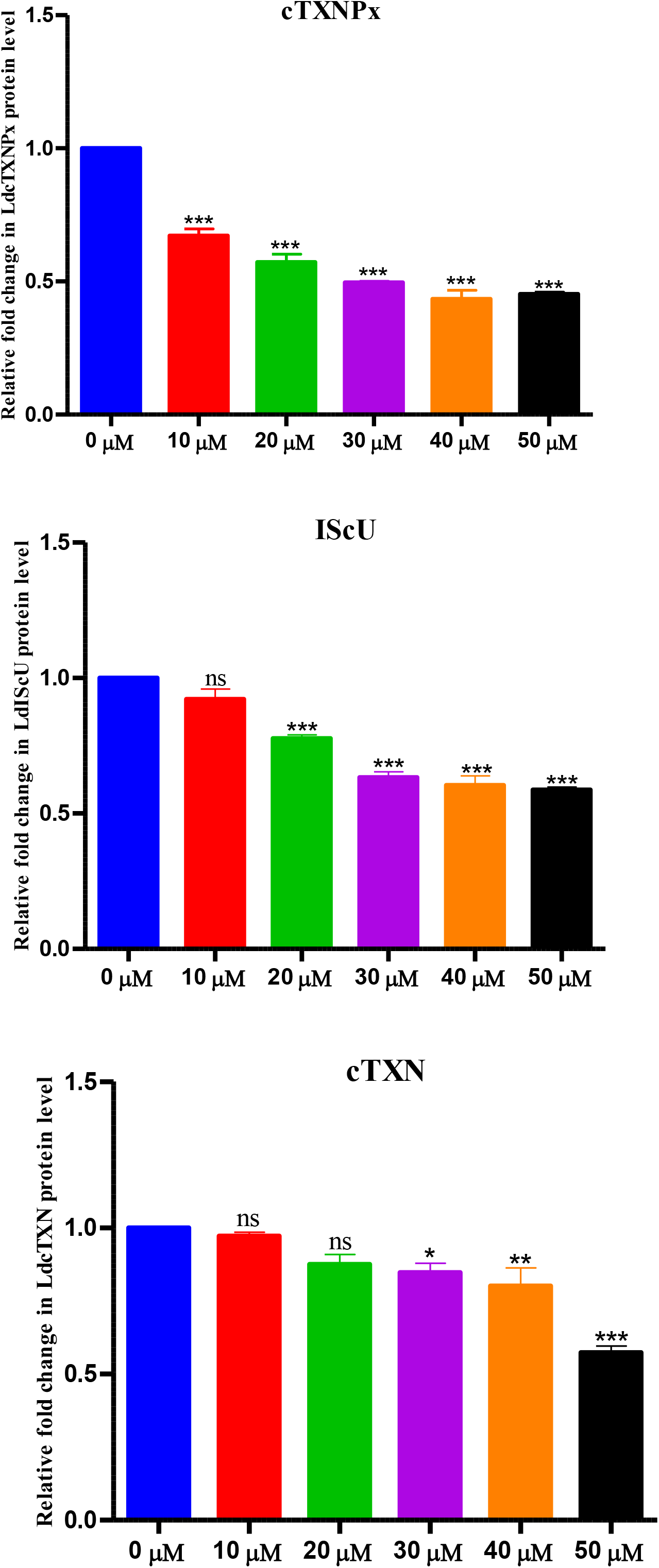
Expression analysis of thiol metabolic and Fe-S clusters proteins and relative fold changes in protein expression of iron depleted promastigote parasites. WT promastigotes were treated with different concentrations of iron-depleted (0-50 µM) conditions for 5 days. After treatment, 30 µg of total cell lysate was loaded on SDS-gel and blotted onto PVDF membrane and further probed with the mentioned *L. donovani* polyclonal antibodies. For loading control, anti-LdActin was used. The western blotting shows expression of *L. donovani* mentioned proteins involved in thiol metabolism and Fe-S cluster biogenesis in iron-depleted conditions. Relative band intensities were determined by performing densitometric analysis using ImageJ software and fold change was calculated by normalising the band intensity with loading control. The abundance of thiol and Fe-S proteins in iron-supplemented **(B)** and iron-depleted conditions **(D)** represented by a graphical plot. Statistical analysis was performed by Student’s t-test and denoted ns for non-significant, *p < 0.05, **p < 0.01, ***p < 0.001 compared to untreated control.

Our findings showed that the lower or higher intracellular availability of iron is important for survival of the parasites and critically involved in modulating the major proteins expression in both thiol and Fe-S cluster biosynthesis pathways in *L. donovani*.

## 4. Discussion

The disease leishmaniasis should be considered as a major health concern globally because it has the potential of massive outbreak. The current management strategy relies on chemotherapeutic drugs, such as liposomal amphotericin B (AmpB), and miltefosine. These drugs have been reported with major drawbacks and reports of regular co-infected cases make it far worse to effectively manage this disease during treatment and post-treatment. For the treatment of PKDL & VL-coinfected cases (HIV-VL, TB-VL) is more complicated because long half-life of miltefosine has been reported for risking resistance emergence. These limitations highlight the urgent need for newer, safer and more effective treatment options by exploring different metabolic pathways in *Leishmania* [2, 42, 43]. Our interest led us to explore how parasites responded at different intracellular iron availability in host cells and utilized their iron during infection and host becomes anaemic which remained poorly explored. However, *Leishmania* completely lacks the ability to synthesize iron thus acquisition and transportation of iron from the host surrounding is critical for the survival of the parasites, both in intracellular (amastigotes) and extracellular (promastigotes) environment [44]. However, with our best knowledge, the precise effect of excess or insufficient intracellular iron on biology in kinetoplastid parasites survival and infectivity remains unaddressed. So, we have showed the first comprehensive analysis of iron supplementation and depletion study that illustrates cell survival, virulence, and redox homeostasis is directly connected to intracellular iron availability, and extremely insufficient iron level leads to cellular stress and apoptosis mediated cell death.

Iron is one such essential micronutrient which regulates most of the biological processes in all living organism [45] including in pathogenic parasites. Iron is an important component for ribonuclotide reductase, which further regulates DNA replication and DNA synthesis. The iron is essential for energy metabolism where iron is required for core proteins, like Fe-S proteins, in the mitochondrial respiration and ATP production as depicted in schematic diagram **(Fig. 14)**. The iron has also been required for heme binding proteins (Cytochrome C, etc) and Fe-S cluster containing proteins, which are involved in the variety of biological events [9, 33, 37, 39]. Loss of iron or decrease in haemoglobin levels lead to anaemia which is considered as a primary symptom during VL patients and overall prevalence has been shown more than 90% [5, 46]. The anaemia has also reported as a major complication in *plasmodium* infection [47].

**Fig 14.**
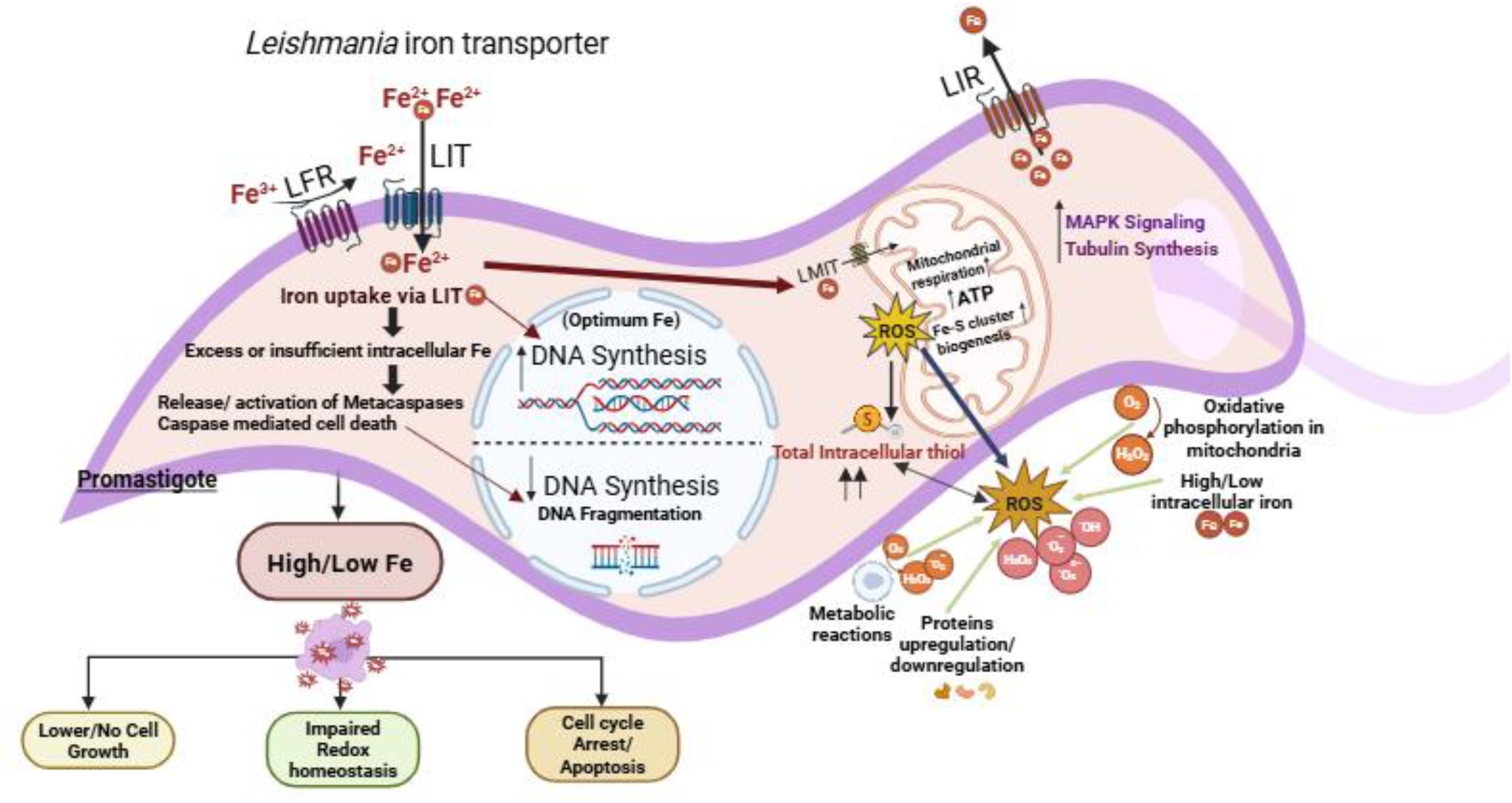
**Proposed model illustrates the function of excess or insufficient iron in promastigotes of *L. donovani***. The schematic representation shows the mechanism of intracellular iron concentration based on our current findings and earlier reports. *Leishmania* utilizes a transmembrane protein *Leishmania* ferric reductase (LFR1) that reduces ferric iron to ferrous form of the iron. Later, the ferrous iron gets acquired and transported inside cell via LIT1. In case of excess or insufficient intracellular iron demand, iron pressure or scarcity creates stress condition, which further triggers release or activation of metacaspases and inhibits DNA synthesis leading to DNA fragmentation and caspase mediated cell death. Optimum intracellular iron is required for promastigote survival and several other metabolic functions. Iron is found critical for cell growth and proliferation, DNA synthesis, mitochondrial function, ATP production, etc. In mitochondria, iron gets transported by a membrane protein known as *Leishmania* mitochondrial iron transporter (LMIT1). Simultaneously, iron has also shown to regulate mitochondrial respiration and Fe-S cluster biogenesis in mitochondria which is responsible for ROS generation. Lower amount of ROS acts as a signalling molecule but during oxidative stress condition ROS production is higher, which drives parasites toward cell growth retardation, and leads to apoptosis. In response to scavange higher level of ROS, intracellular thiol is increased to maintain redox balance. Intracellular ROS level can be accelerated by other cellular events like oxidative phosphorylation in mitochondria, intracellular iron stress, metabolic reactions and protein pathways upregulations or downregulation. Besides, increased ROS causes oxidative damage to flagellar protein which showed flagellar shortening in parasite morphology in both high and low iron condition. However, these parasites have also known to have *Leishmania* iron regulator (LIR), another transmembrane protein, which regulates their cytosolic accumulated of iron to prevent intracellular iron toxicity. This LIR1 acts as exporter and shuttles iron outside of the cell but somehow this LIR1 protein gets switch off in excess or insufficient iron concentration, which is poorly explored. Furthermore, we have shown that iron also causes morphological changes and literature findings suggested that iron also effects cytoskeletal remodelling, tubulin synthesis and MAP kinase which drives, promastigote elongation and metacyclogenesis. This diagram is created with biorender.com.

Apart from iron, other metals have also been explored in different parasite biology. Previous studied have too reported that other metals like Zn, Cu, Ca, Mn & Se are important in parasite biology, acting as cofactors, modulating virulence, and providing structural integrity [48–50]. The disruption of metal homeostasis undermines *Leishmania* promastigote morphology, size, and flagellar structure. While essential metals like Fe, Zn, Cu, Mg & Ca support normal cellular functions, their imbalance leads to structural and functional impairments [51]. The studies have showed that iron is essential for survival and regulation of many biochemical processes including mitochondrial metabolism, parasite growth and proliferation, oxidative stress management, parasite virulence, and disease progression in *Leishmania* [35, 39, 52–54] *Plasmodium* [55] and *Trypanosomatids* [8, 56, 57]. *Leishmania* uses distinct iron acquisition mechanism including *Leishmania* Ferric Reductase (LFR1) [11, 17] and *Leishmania* iron transporter (LIT1) [10, 13, 53], both of which are located on the plasma membrane and mediate iron import in the parasitophorous vacuoles (PV). Another, plasma membrane protein *Leishmania* Iron Regulator (LIR1), is an intracellular iron exporter which regulated iron homeostasis and promotes infectivity [36, 58]. Iron has also found significant for plants and vacuolar iron transporter (VIT) has already been reported in different organism including plants for iron transportation [59, 60]. Similaly, zinc showed a crucial role in cell survival and its deficiency led to mitochondrial dysfunction leading to apoptosis like cell death in *Leishmania* [19, 27] and *Plasmodium* [61]. While, zinc supplementation led to more efficient immune response reducing the pathogenic effect in *Trypanosoma* infection [62]. Copper is reported significant for cytochrome oxidase and superoxide dismutase (SOD) activity, but in excess induces oxidative damage, leading to cell shrinkage and flagellar loss. Copper was also found significant in oxidative stress defense, pathogenicity and successful manifestation of infection in *Leishmania* [63] and *Trypanosoma* [64]. Calcium is reported to regulate flagellar motility and wave polarity in *L. donovani* [65], meanwhile, Manganese (Mn) and selenium (Se) also reported to contribute in parasite survival by detoxifying reactive oxygen species in *Schistosoma* [66]. Despite all other metals, iron has shown to play more vital role in parasite survivability and infectivity [10, 11, 13, 17, 36, 53, 58]. Our present study demonstrates the impact of intracellular iron supplementation (ferrous sulphate) and iron depletion (deferoxamine) on the growth, viability, and pathogenicity of *L. donovani* promastigotes. Recently, an interesting study has been reported which deciphered the mechanism of iron acquisition/utilization by focusing on iron transportation that facilitated parasite *L. donovani* resistant strain rapid proliferation. This work also revealed major molecular events leading to severe anaemia in response to *L. donovani* infected mice macrophages [15]. In this present study, we have examined the impact of low or higher iron concentrations in *L. donovani,* where we found the parasite survival is directly proportional to intracellular iron level. Parasites always require optimum iron concentrations for better survivability and growth. Our result showed iron acts as a metabolic regulator in *L. donovani* and parasite thrives in iron-rich environment **(Fig. 1B)** but iron deprivation *via* DFO mimics antimicrobial defense slowing parasite survival **(Fig. 2B)**. The iron estimation revealed intracellular iron was increasing with iron supplementation **(Fig. 4A)**. Iron chelation by DFO led to iron deprived condition as showed by intracellular iron estimation **(Fig. 4B),** where parasite showed very restricted growth with increasing iron chelation. Therefore, it is evident that Fe is essential for the growth of *L. donovani* parasites.

Further, morphological analysis showed that iron supplementation led to elongated promastigotes cell shape and longer flagella with higher motility, possibly due to iron’s role in metabolic enzymes and cytoskeletal remodelling. Excess iron can induce oxidative stress, causing structural deformities such as irregular cell shapes and reduced motility. Conversely, iron restriction reduces promastigote growth and leads to shorter, rounded cells with stunted flagella indicative of cellular stress and impaired motility. These changes indicate that iron is necessary for flagellar biogenesis and its defects impair motility, crucial for host invasion. Our result revealed intracellular iron also impacts parasite morphology **(Fig. 5A & B).** By considering our result and previous studies, we concluded that iron regulates tubulin synthesis and involved in cellular microtubule elongation. Also, the Mitogen-activated protein (MAP) kinases signalling have been reported earlier shown to drive promastigote elongation and flagellar length control in *Leishmania* [67]. We also observed initial increase (0-500 µM) and then reduced (1000 & 2000 µM) formazan formation in MTT assay in iron supplemented condition, which serves as a marker of cell viability while iron depletion showed reduction in formazan formation and exhibited lower cell viability (**Fig 6)**. Recently, the association of gp63, a Zn dependent metalloprotease has been shown associated with iron regulation [68]. These gp63 are involved in host-parasite interaction and virulence reported earlier [69]. To establish the physiological relevance of iron-mediated effects on infectivity, we investigated the effect of iron in *in vitro* infection studies using THP-1 derived macrophages **(Fig. 7A & 8A)**. Iron supplementation was found to enhance *L. donovani* promastigotes infectivity **(Fig. 7B)**, whereas iron depletion significantly reduces parasite infectivity **(Fig. 8B)**. These findings demonstrate that iron is important in promoting parasite survival and differentiation within of host macrophages also documented in other earlier study for zinc [19, 70]. Iron supplementation has revealed higher infectivity compared to other divalent metals, indicating a prominent role of iron in parasite virulence and infectivity. Additionally, similar metal-dependency has been reported for zinc in another study, where depletion impaired infectivity by disrupting redox systems and mitochondrial stability [27].

These results showed the importance of iron in cell morphology and viability. Iron can catalyse the formation of reactive oxygen species (ROS) through Fenton’s chemistry, leading to massive oxidative stress and cellular damage [38]. Our findings revealed that lower iron concentrations (0-1000 µM) promote parasite growth and proliferation; while higher concentrations are toxic (2000 & 4000 µM) resulted in lower cell proliferation also reported earlier in zinc and another study for LIR1 protein in *Leishmania* [19, 36]. By using ferrous sulphate and deferoxamine, we detected involvement of caspase or caspase like activity in *L. donovani* cell death depending upon iron concentration. The caspase like activity mediated cell death was also showed in zinc as reported earlier [71]. We also detected substantial increase with significant fold change in caspase like activity in both 1000 & 2000 µM iron supplementation and 30 & 50 µM depletion condition. Notably, higher concentrations of both iron supplementation and depletion resulted in increased cellular stress leading to caspase-mediated apoptosis like cell death and enhanced DNA fragmentation, suggesting a direct impact of iron on parasite survival **(Fig. 9A & 9B)**. The intracellular iron concentration driven caspase-like activity, DNA fragmentation and apoptosis has shown first time in present study which magnifies the importance of iron in cellular events. However, apoptosis has been already reported as one of the major event in Leishmanial parasites death [72, 73].

However, till date no reports related to exposure of concentration dependent iron depletion and iron supplementation to *L. donovani* parasites, showed impact on alterations in infectivity, Fe-S and thiol protein expression, caspase-like activity, DNA fragmentation and apoptosis has been reported so far best of my knowledge. Other molecular cell death mechanism pathways has been proposed in protozoan cell death through necrosis and incidental death [74] and both caspase-dependent and caspase-independent mechanism for *Leishmania spp.* apoptosis are shown in different studies [75–77]. However, we concluded high iron supplementation in parasites (1000 & 2000 µM) possibly increased the intracellular Fe^2+^ labile pool resulted in release of metacaspases and nucleases led to proteolysis with activation of endonucleases as reported in other study [78]. This resulted in nucleosomal DNA fragmentation as shown in results **(Fig. 9C & 9D).** While low intracellular iron impaired Fe-S clusters assembly leading to massive oxidative stress, which similarly resulted in caspase like activity mediated cell death. With DFO, our result revealed higher DNA fragmentation as illustrated in schematic diagram **(Fig. 14)**.

We have further evaluated ROS generation and total intracellular thiol content, both of which serve as indicators of oxidative stress and antioxidant capacity. ROS has known as a major cause for growth retardation and cell death [79]. ROS determines cellular fate as it has considered as a significant signalling molecule which further regulates several proteins and important cellular components [80]. To validate the reason behind reduced parasite growth and viability, we found that a significant flux in intracellular ROS in both iron supplemented and iron-depleted conditions in comparison to control parasites. However, certain nutrient deprivation condition has been previously reported to induce ROS in *Leishmania* parasites [81]. Our research findings demonstrated that iron supplementation enhances parasite tolerance toward external oxidative stress tolerance of iron by reducing intracellular reactive oxygen species (ROS) accumulation in parasites and increased total intracellular thiol content led to parasite survival even at higher iron supplementation with increasing time, suggesting upregulation of antioxidant pathways including non-protein thiols like trypanothione-a hallmark of parasite redox defense in wildtype parasites similarly reported earlier [82]. In contrast, iron depletion parasites lead to increased oxidative damage, evidenced by higher ROS levels, and decreases in total intracellular thiol content in *L. donovani* parasites **(Fig 10)**. These changes may disrupt essential metabolic functions within the parasites. Thus, these observations prompted us to study the expression pattern of different Fe-S and thiol metabolic pathway proteins as we have shown. Our findings suggest that adequate iron levels are vital for maintaining parasite redox homeostasis and metabolic stability, while iron deficiency compromises their ability to manage oxidative stress [15, 57, 83]. Our result also revealed that iron supplemented parasites were able to tolerate additional oxidative stress generated by 2.5 µM and 5.0 µM menadione treatment. While iron depleted parasite were not able to cope with menadione stress and cell death because of excess oxidative stress where growth was completely retarded **(Fig. 11)**.

To better understand, the molecular basis of response, we have further assessed *L. donovani* Fe-S cluster assembly proteins (Nbp35, Frataxin & IScU) which are essential for mitochondrial function, electron transport and DNA metabolism as well as thiols associated antioxidant proteins (TryS, TryR, CS, SAT, cTXN, & cTXNPx) expression under varying iron supplemented (0-4000 µM) and iron depleted (0-50 µM) conditions. In addition, our result showed iron effects thiol metabolism proteins as result showed in iron supplementation and depletion. Iron not only effects iron related proteins but also effects thiol containing proteins which were either upregulated or downregulated as shown in results (**Fig. 12 & 13)**. This showed the inter-connection between iron and thiol metabolism as reported earlier [84]. Our findings revealed that most of the thiol and Fe-S pathway proteins were upregulated in response to iron supplementation and showed significant fold change at higher iron concentration suggesting that iron enhances mitochondrial activity and redox maintenance mechanisms, as Fe-S clusters biosynthesis is iron-dependent and central to severe redox sensitive enzymes and adequate iron levels are essential for the proper functioning and assembly of Fe-S clusters as well as the synthesis and maintenance of thiol-based antioxidant systems. The enhanced expression under iron-rich conditions likely reflects the parasite’s adaptive mechanism to utilize available iron for vital metabolic processes, thereby supporting redox homeostasis and survival. Interestingly, under iron depleted conditions, the expression of most proteins were downregulated and showed significant lower fold change, resulted in the impaired metabolic activity expected during iron starvation in parasites. However, Frataxin, a mitochondrial iron binding/carrier protein and regulator, was upregulated under challenged iron-deprived conditions suggested intracellular iron scarcity utilized mitochondrial iron storage protein Frataxin as a coping mechanism for fulfil of iron demand to meet survival. This contrasts with other Fe-S cluster proteins and may represent a protective mechanism to buffer intracellular iron stress, as seen in [85], which highlight Frataxin’s role in iron detoxification and storage under stress conditions.

Therefore, iron supplementation stimulates parasites proliferation by enhancing energy metabolism & DNA synthesis while DFO inhibits the growth by inducing iron starvation stress, ROS imbalance and mitochondrial dysfunction **(Fig. 14).** The iron supplementation parasites shows higher infection which suggests iron is required by amastigotes for growth and infection in parasitophorous vacuoles of macrophages. This also highlights that in VL patients, the amastigotes sequest iron from host led to anaemia development during VL. Together, these findings demonstrated that adequate iron availability appears to facilitate essential metabolic and redox processes necessary for parasite establishment, while iron deficiency impairs these functions, leading to diminished infectivity. These findings underscore the delicate balance between *Leishmania* and iron concentrations that must be maintained to regulate iron concentrations to avoid cytotoxicity. The toxic effects of both iron excess and deficiency suggest that tightly regulated iron concentration is essential for *Leishmania* survival. This underscores the importance of iron homeostasis in the host-parasite interaction and suggests that modulating iron levels could influence the course of infection. Altogether, this work paves the way for future studies to explore the role and significance of iron related pathway in visceral leishmaniasis.

## Conclusion

Our study highlights that iron is essential for the parasite survival, cellular redox homeostasis, and pathogenicity of *L. donovani.* Our findings suggest that excess or insufficient iron acquisition led to cytotoxicity, further emphasizing the importance of tightly regulated iron homeostasis. Our result showed intracellular iron supplementation and depletion significantly alters Fe-S and thiol metabolism proteins expressions. We further explored the role of iron in ROS and intracellular thiol stress management which effectively triggered caspase-mediated cell death pathway in *Leishmania* parasites. This study also revealed that parasite virulence is directly related to iron availability and the iron-rich condition as showed increased percentage infectivity because high iron load led to higher infection in THP-1 macrophages. Our study emphasizes that iron concentration may lead anaemic prevalence in VL patients as parasites required iron for growth may be used iron form host causing disruption of RBCs and haemoglobin or other iron binding proteins. Collectively, these results highlight the importance of iron in parasite growth, differentiation, redox imbalance, oxidative stress, protein upregulation and downregulation, iron influenced cell death and macrophage infectivity.

## Acknowledgement

The authors would like to thank ICMR-RM NIHR, Patna and the Indian Council of Medical Research, New Delhi for providing the research facilities and funding assistance through an intramural ICMR project to VA. This work was supported with the grant no. (INT-170-BAS/2020) by ICMR. This study was also funded by ICMR [Grant No. 6/9-7 (259)/KA/2021/ECD-II], ICMR-RMRIMS (INT-135-BAS/2017), Department of Health Research and Department of Science and Technology [Grant No. DST/CCP/NHH/113/2017(G)] given to VA. Dr. Amogh Sahasrabudhhe from CSIR-CDRI, Lucknow is sincerely acknowledged for providing β-actin antibody for this study. PP expresses her profound gratitude to the University of Calcutta for supporting with Ph.D. programme. PP is also thankful to ICMR for providing ICMR-SRF fellowship support during Ph.D. tenure with letter no. Fellowship/94/2022-ECD-II. PP deeply acknowledges Mr. Vimal Kumar for providing camera to picture macrophage infectivity experiment. PP is grateful for lab members and the technical staff for their help, co-operation and support.

## Declaration of competing interest

The authors declare no conflicts of interest. The authors report no financial or personal relationships that could have influenced this work.

---

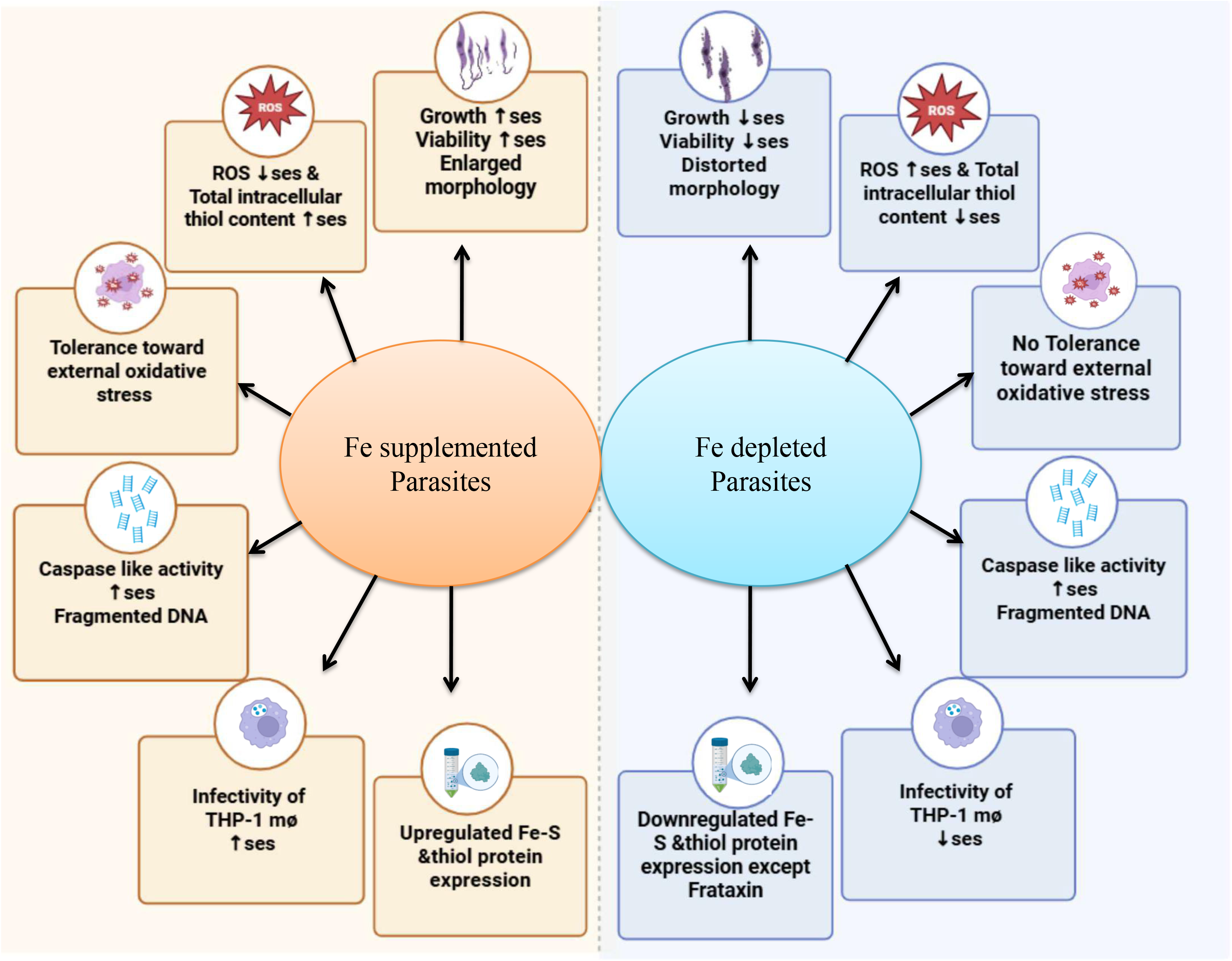

